# A systematic assessment of phylogenomic approaches for microbial species tree reconstruction

**DOI:** 10.1101/2024.11.20.624597

**Authors:** Samson Weiner, Yutian Feng, J. Peter Gogarten, Mukul S. Bansal

## Abstract

A key challenge in microbial phylogenomics is that microbial gene families are often affected by extensive horizontal gene transfer (HGT). As a result, most existing methods for microbial phylogenomics can only make use of a small subset of the gene families present in the microbial genomes under consideration, potentially biasing their results and affecting their accuracy. To address this challenge, several methods have recently been developed for inferring microbial species trees from genome-scale datasets of gene families affected by evolutionary events such as HGT, gene duplication, and gene loss.

In this work, we use extensive simulated and real biological datasets to systematically assess the accuracies of four recently developed methods for microbial phylogenomics, SpeciesRax, ASTRAL-Pro 2, PhyloGTP, and AleRax, under a range of different conditions. Our analysis reveals important insights into the relative performance of these methods on datasets with different characteristics, identifies shared weaknesses when analyzing complex biological datasets, and demonstrates the importance of accounting for gene tree inference error/uncertainty for improved species tree reconstruction. Among other results, we find that (i) AleRax, the only method that explicitly accounts for gene tree inference error/uncertainty, shows the best species tree reconstruction accuracy among all tested methods, (ii) PhyloGTP (developed previously by the authors of this paper) shows the best overall accuracy among methods that do not account for gene tree error and uncertainty, (iii) ASTRAL-Pro 2 is less accurate than the other methods across nearly all tested conditions, and (iv) explicitly accounting for gene tree inference error/uncertainty can lead to substantial improvements in species tree reconstruction accuracy. Importantly, we also find that all methods, including AleRax and PhyloGTP, are susceptible to biases present in complex real biological datasets and can sometimes yield misleading phylogenies.

## 1 Introduction

The accurate inference of phylogenetic relationships between different microbes is an important problem in evolutionary biology. A key difficulty in estimating such phylogenies is the presence of extensive horizontal gene transfer (HGT) in microbial evolutionary histories [32]. This can result in markedly different evolutionary histories for different gene families, obfuscating the underlying species-level or strain-level phylogeny. As a result, the traditional approach for reconstructing microbial phylogenies is to use only “well-behaved” gene families resistant to HGT. This includes the use of small-subunit ribosomal RNA genes (e.g., [47, 68]) or of a concatenated alignment of a few core genes from the genomes of interest (e.g., [11, 31, 38]). Both these approaches, however, are known to be error-prone. For instance, ribosomal RNA genes are known to engage in horizontal transfer [21, 69, 71] and to yield histories that are inconsistent with those inferred using other core genes [15, 16, 24, 25]. Furthermore, ribosomal RNA genes often cannot be used when studying closely related species due to excessive sequence similarity. Similarly, concatenation based approaches, such as the widely used multilocus sequence analysis (MLSA) technique [20], essentially ignore HGT and aggregate the phylogenetic signal from several gene families with potentially distinct evolutionary histories [19, 40]. Indeed, the tree resulting from the concatenation might represent neither the organismal phylogeny nor any of the genes included in the concatenation [34].

To overcome these limitations, several genome-scale methods have also been proposed for microbial phylogeny inference. These include methods such as Phylo SI that are based on gene order information [55, 56], supertree-based methods such as SPR supertrees [65] and MRP [6, 71] that allow for the use of multiple orthologous gene families, and methods based on average nucleotide identity (ANI) of genomes [22, 23, 28]. Such genome-scale methods are inherently preferable to methods that base phylogeny reconstruction on only a single gene or a small set of concatenated genes [40]. However, while these above methods all represent useful approaches for microbial phylogenomics, they are either targeted at analyzing closely related strains or species (gene order and ANI based methods), or are limited to using single-copy gene families or orthologous groups and do not model key evolutionary events affecting microbial gene family evolution (supertree based methods). Recently, truly genome-scale approaches for microbial phylogenomics, capable of using thousands of complete (multi-copy) gene families, have also been developed. Four of the most promising such methods are ASTRAL-Pro 2 [70], SpeciesRax [44], PhyloGTP [64], and AleRax [45]. These methods all take as input a collection of unrooted gene trees, where each gene tree may contain zero, one, or multiple genes from a species/strain under consideration. ASTRAL-Pro 2 is based on quartets and seeks a species tree that maximizes a quartet based score [70]. While ASTRAL-Pro 2 does not directly model any specific evolutionary processes, such as HGT or gene duplication, responsible for gene tree discordance, it can handle complete (multi-copy) gene families and previous research suggests that it’s quartet based approach should be robust to HGT [13]. SpeciesRax uses an explicit Duplication-Transfer-Loss (DTL) model of gene family evolution in microbes and seeks a species tree that maximizes the reconciliation likelihood of observing the input gene trees under that model [44]. PhyloGTP, a method previously developed by the authors of the current paper, takes a similar overall approach as SpeciesRax but is based on the gene tree parsimony approach and uses a different heuristic search strategy. Specifically, PhyloGTP uses a parsimony-based DTL framework to account for HGT, gene duplication, and gene loss and uses local search heuristics to find a species tree with lowest total reconciliation cost with the input gene trees [64]. AleRax performs species tree inference under a probabilistic DTL model [45] and is more sophisticated than the other methods in that it can co-estimate gene trees along with the species tree. Unlike ASTRAL-Pro 2, SpeciesRax, and PhyloGTP, which all take as input a single, fixed gene tree per gene family, AleRax takes as input multiple MCMC gene tree samples for each gene family and can thus explicitly account for gene tree reconstruction error and uncertainty.

In this work, we use an extensive simulation study and two real biological datasets to evaluate the species tree reconstruction accuracies of ASTRAL-Pro 2, SpeciesRax, PhyloGTP, and AleRax. Our simulation study focuses on systematically evaluating the impact of number of input gene trees, realistic rates of duplication, HGT, and loss events, and input gene tree error rates on all methods, and on evaluating the impact of using multiple gene tree samples on AleRax. We find that AleRax, the only method that explicitly accounts for gene tree inference error/uncertainty, shows the best species tree reconstruction accuracy among all tested methods, while PhyloGTP shows the best overall accuracy among methods that do not explicitly account for gene tree error and uncertainty. AleRax shows similar accuracy as PhyloGTP when using true (error-free) simulated gene trees, but yields a substantial improvement in reconstruction accuracy compared to PhyloGTP when using estimated (error-prone) simulated gene trees. Between PhyloGTP and SpeciesRax, we find that PhyloGTP can substantially outperform SpeciesRax when the number of input gene trees is small or when DTL rates are high, while SpeciesRax often outperforms PhyloGTP on datasets with low DTL rates. ASTRAL-Pro 2 shows worse accuracy than all other methods across nearly all tested conditions. Overall, the average reconstruction accuracies (defined formally later) of ASTRAL-Pro 2, SpeciesRax, PhyloGTP, and AleRax across our core simulated datasets with estimated gene trees are 81.2%, 86.9%, 88.8% and 91.3%, respectively. We find, however, that the improved accuracies of AleRax and PhyloGTP over the other methods come at the expense of substantially longer running times. We also investigate how AleRax’s ability to handle gene tree error/uncertainty contributes to its species tree reconstruction accuracy. We find that when AleRax is provided as input only a single estimated gene tree per gene family (as with the other methods), its accuracy becomes comparable to that of PhyloGTP. This suggests that explicit handling of gene tree error/uncertainty can lead to an approximately 20% reduction in species tree reconstruction error.

We also use the four methods to analyze two real microbial datasets; a more complex 174-taxon Archaeal dataset exhibiting extreme divergence and compositional biases, and a less complex dataset of 44 Frankiales exhibiting low divergence. We find that all four methods perform well on the less complex dataset, recovering identical relationships among the major clades. On the more complex dataset, PhyloGTP, SpeciesRax and ASTRAL-Pro 2 result in some incorrect placements, but appear to perform better than AleRax which produces a tree that is markedly different than any highly supported previously calculated Archaeal tree. Overall, this suggests that all tested methods, including AleRax and PhyloGTP, are potentially susceptible to biases present in complex datasets.

Overall, our results indicate that AleRax and PhyloGTP may be the two best methods currently available for microbial phylogenomics, though their improved accuracies come at the cost of significantly longer running times. Our results also suggest that phylogenomics methods can benefit substantially from explicitly account for gene tree error and uncertainty. At the same time, our results show that even the best existing methods for microbial phylogenomics may not produce accurate results for certain complex microbial datasets and that their results should be interpreted with caution. A preliminary version of this manuscript, which focused on describing and evaluating PhyloGTP, appeared in the proceedings of the RECOMB Comparative Genomics 2024 conference [64]. The current manuscript focuses more heavily on a systematic experimental assessment of the four methods and expands upon the preliminary version by (i) including the recently published method AleRax in the experimental evaluation, (ii) studying the impact of event cost assignments on PhyloGTP’s accuracy, (iii) providing descriptions of all four methods evaluated, (iv) evaluating the methods on additional datasets with different ratios of evolutionary events, (v) evaluating memory requirements of all methods, (vi) providing an updated and expanded assessment of the methods on the two real datasets, and (vii) presenting a more extensive discussion of the experimental evaluation and our findings.

## 2 Materials and Methods

### 2.1 Description of Evaluated Methods

We provide brief descriptions of the four methods considered in this work and state their specific objective functions.

**Basic definitions and preliminaries.** Let *T* be a leaf-labeled tree with node, edge, and leaf sets denoted by *V* (*T*), *E*(*T*), and *Le*(*T*). If *T* is rooted, we denote its root by *rt*(*T*). For any node *v ∈ V* (*T*), where *T* is a rooted tree, the (maximal) subtree rooted at *v* is denoted *T_v_*. Unless otherwise specified, all trees are binary and unrooted.

We use the term *species tree* for the tree depicting evolutionary relationships for the taxa (e.g., species, strains, etc.) under consideration. Given a gene family from the taxa under consideration, a *gene tree* is a tree that depicts the evolutionary relationships of the genes in the gene family. We assume that each edge in a gene tree has an associated branch length (representing substitutions per site), though not all methods make use of branch lengths. Note that a gene tree may have zero, one, or multiple genes from the same taxon.

We assume that the taxon set under consideration is denoted by Ω and that the species tree, denoted *S*, depicts the evolutionary relationships for taxa in Ω, i.e., *Le*(*S*) = Ω. We use *G* to denote a collection of gene trees *{G*_1_*, …, G_k_}*, where each *G_i_*, 1 *≤ i ≤ k*, describes the evolutionary history of a gene family present in the taxon set Ω. We implicitly assume that *Le*(*S*) = *∪^k^ Le*(*G_i_*). ASTRAL-Pro 2, SpeciesRax, and PhyloGTP assume that each input gene tree (i.e., each gene tree in *G*) corresponds to a different gene family, while AleRax requires multiple gene tree samples per gene family, as explained below. All methods considered in this work assume that the input gene trees are unrooted.

The methods SpeciesRax, AleRax, and PhyloGTP utilize DTL reconciliation to assess the fit of input gene trees with candidate species trees. DTL reconciliation provides a framework for reconciling the differences between a gene tree and the corresponding rooted species tree by invoking gene duplication, HGT, and gene loss events. The method ASTRAL-Pro 2 uses quartet trees to assess the fit between the input gene trees and candidate species trees. A quartet tree is an unrooted tree on four leaves, and the number (or fraction) of quartet trees shared between a gene tree and an unrooted species tree can serve as a measure of similarity between the two trees. Further details on each method appear below.

**ASTRAL-Pro 2.** This quartet-based method builds upon the widely used ASTRAL method [42]. ASTRAL is designed to work with single-copy gene trees constructed from orthologous sequences and seeks a species tree that maximizes the *quartet score* with the input gene trees. ASTRAL-Pro 2 uses a related but different similarity measure called *per-locus quartet score*, designed to avoid over-counting of quartets in multi-copy gene trees. Thus, given a collection of unrooted gene trees *G* as input, ASTRAL-Pro 2 seeks an unrooted species tree *S* that maximizes the per-locus quartet score with the input gene trees. Since the underlying computational problem is NP-hard [30], ASTRAL-Pro 2 implements a heuristic for the problem, using dynamic programming to efficiently find an optimal species tree within a restricted search space. We note that ASTRAL-Pro 2 does not make use of gene tree branch lengths. Further details on the method appear in [70].

**SpeciesRax.** This method uses the “undatedDTL” probabilistic DTL reconciliation framework of GeneRax [43] to estimate the species tree and model parameters (rates of duplication, HGT, and loss) given a collection of input gene trees. Specifically, SpeciesRax takes a collection of unrooted gene trees *G* as input and seeks a rooted species tree *S* and model parameters Θ that maximize the reconciliation likelihood *L*(*S,* Θ*|G*). SpeciesRax uses a distance-based method, *MiniNJ*, to estimate a starting species tree and then executes a local search heuristic to further optimize this starting tree. We note that SpeciesRax utilizes gene tree branch lengths and infers a *rooted* species tree. Further technical details on SpeciesRax appear in [44].

**AleRax.** AleRax is similar to SpeciesRax in that it also uses a probabilistic model of DTL reconciliation and uses the same search strategy as SpeciesRax (miniNJ followed by local search) for finding a maximum likelihood species tree and model parameters given the input gene trees. However, unlike SpeciesRax, AleRax accounts for gene tree inference error and uncertainty by taking as input multiple gene tree samples for each gene family. Specifically, AleRax uses the ALE algorithm [59] to integrate over gene tree uncertainty by approximating the probability of observing a gene family sequence alignment given a rooted species tree. Thus, AleRax seeks to find a rooted species tree *S* and model parameters that maximize the likelihood *L*(*S|A*), where *A* denotes the collection of sequence alignments for all gene families represented in *G*. We note that AleRax does not directly take sequence alignments as input and instead uses the multiple (typically 1000) input gene tree samples for each gene family sequence alignment to estimate the fit of an alignment with a species tree. Like SpeciesRax, AleRax utilizes gene tree branch lengths and infers a *rooted* species tree. Further details on this method appear in [45].

**PhyloGTP.** Unlike SpeciesRax and AleRax, PhyloGTP uses the parsimony-based DTL reconciliation model first developed in [3, 12, 60]. Under this model, each event type has an associated (user-defined) cost and the objective is to find a reconciliation of minimum total cost. This model allows for an unrooted gene tree to be optimally reconciled with a rooted species tree within *O*(*mn*) time, where *m* and *n* denote the number of leaves in the gene tree and species tree, respectively [3].

In the following, we denote the event costs for gene duplications, HGTs, and gene losses by *P_d_*, *P_t_*, and *P_l_*, respectively. Given a gene tree *G ∈ G*, species tree *S*, and event costs *P_d_*, *P_t_*, and *P_l_*, we denote by *R_Pd,Pt,Pl_* (*G, S*) the reconciliation cost of an optimal DTL reconciliation of *G* and *S* under the event costs *P_d_*, *P_t_*, and *P_l_*. Given a species tree *S*, a collection of gene trees *G* = *{G*_1_*, …, G_k_}*, and event costs *P_d_*, *P_t_*, and *P_l_*, we define the *total DTL reconciliation cost* of *G* with *S* to be the sum of the DTL reconciliation costs of each *G ∈ G* with *S*, i.e., 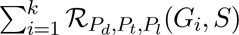.

Given as input a collection of gene trees, PhyloGTP seeks a species tree that minimizes the total DTL reconciliation cost against the collection of input gene trees. More formally, we can define the *Most Parsimonious Species Tree (MPST)* problem as follows: Given a collection of gene trees *G* and event costs *P_d_*, *P_t_*, and *P_l_*, find a species tree *S* that minimizes the total DTL reconciliation cost with *G*.

We note that under this formulation only the topology of the gene tree is used and branch lengths are ignored. The MPST problem can be shown to be NP-hard, W[2]-hard, and inapproximable to within log factor through a reduction from the NP-hard gene duplication problem [4, 36]. The gene duplication problem is a special case of MPST problem defined in this manuscript and seeks a species tree minimizing just the total number of gene duplications. Details of the reduction are straightforward and therefore omitted. PhyloGTP uses a local search heuristic to solve the MPST problem. For completeness, further methodological details on PhyloGTP appear in the Supplement.

### 2.2 Experimental setup

We use both simulated and real biological datasets to carefully assess the reconstruction accuracy of the four methods. ASTRAL-Pro 2, SpeciesRax, and PhyloGTP take as input a single unrooted maximum-likelihood gene tree per gene family, while the recommended input for AleRax is 1000 unrooted gene trees, sampled from the posterior using Mr.Bayes [54], per gene family. All methods were run using their default/recommended parameter settings. For AleRax, we present results both for the recommended number (1000) of gene trees per gene family, allowing it to account for gene tree inference error/uncertainty, as well as when only a single gene tree is used per gene family, effectively disabling its ability to account for gene tree error/uncertainty.

**Evaluating reconstruction accuracy.** To evaluate the species tree reconstruction accuracies of the different methods, we compare the species tree estimated by each method with the corresponding ground truth species tree. To perform this comparison we utilize the widely used (unrooted) normalized Robinson-Foulds distance (NRFD) [53] between the reconstructed and ground truth species trees. For any reconstructed species tree, the NRFD reports the fraction of non-trivial splits in that species tree that do not appear in the corresponding ground truth species tree. For ease of interpretation, we report results in terms of *percentage accuracy*, defined to be the percentage of non-trivial splits in the reconstructed species tree that also appear in the ground truth species tree. Thus, percent accuracy is simply (1 *−* NRFD) *×* 100. Thus, for example, a percentage accuracy of 87% is equivalent to an NRFD of 0.13.

### 2.3 Description of simulated datasets

We used simulated datasets with known ground truth species trees to assess the impact of three key parameters on reconstruction accuracy: Number of input gene trees, rates of gene duplication, HGT, and gene loss (or DTL rates for short), and estimation error in the input gene trees.

Our core simulated datasets were created using a four-step pipeline: (1) simulation of a ground-truth species tree and corresponding true gene trees (one per gene family) with varying DTL rates, (2) simulation of sequence alignments of different lengths for each true gene tree, (3) inference of estimated maximum likelihood gene trees from the sequence alignments, and (4) inference of 1000 estimated gene trees, sampled from the posterior using Mr.Bayes [54], for each true gene tree (i.e., per gene family) from the sequence alignments. In the first step, we used SaGePhy [29] to first simulate ground-truth species trees, each with exactly 50 leaves (taxa) and a height (root to tip distance) of 1, under a probabilistic birth-death framework. We then used these species trees to simulate multiple gene trees under the probabilistic duplication-transfer-loss model implemented in SaGePhy. This resulted in 9 different datasets of simulated true gene trees, each corresponding to a different number of true input gene trees (10, 100, or 1000), and a different DTL rate (low, medium, or high; see Table 1). Each dataset comprised of 10 replicates. The chosen DTL rates are based on the relative rates and frequencies of gene duplication and HGT events in real microbial gene families from species sampled broadly across the tree of life [5, 12]. In each case, the gene loss rate is assigned to be equal to the gene duplication rate plus the additive HGT rate, so as to balance the number of gene gains with the number of gene losses (Table 1). Basic statistics on these simulated true gene trees, including average sizes and numbers of gene duplication and HGT events, are provided in Table 2. We note that numbers of inferred gene duplications and HGTs are larger for estimated gene trees, where reconstruction error manifests itself as closely matching rates inferred for real microbial gene families based on estimated gene trees [5, 12].

**Table 1.** Key parameters used to generate the core simulated datasets. The table lists the main parameters and their values explored in the simulation study for the core datasets. All 36 (= 3 *×* 3 *×* 4) combinations of these three parameters were evaluated at 10 replicates each. DTL rates are specified in the form (*d, t, l*), where *d* is the gene duplication rate, *t* is the HGT rate (split evenly between additive and replacing HGTs), and *l* is the gene loss rate. The number of species was fixed at 50 for these datasets.

| Parameter | Values |
| --- | --- |
| Number of gene families | 10, 100, 1000 |
| DTL rates | low = (0.3, 0.6, 0.6)<br>med = (0.6, 0.12, 0.12)<br>high = (0.12, 0.24, 0.24) |
| Sequence length (nucleotides) | 400, 100, 50, and true gene trees |

**Table 2.** Basic statistics for simulated true gene trees in the core dataset. Average number of leaves, duplications, and HGTs, and losses in the simulated low, medium, and high DTL true gene trees in the core simulated datasets. For each DTL rate, the number of losses is roughly equal to the number of duplications plus half the number of HGTs. Results were averaged over all 10 replicates of the 100 gene tree datasets.

| DTL rate | Leaves | Duplications | HGTs |
| --- | --- | --- | --- |
| Low | 53.618 | 3.408 | 6.586 |
| Med | 55.121 | 6.15 | 11.125 |
| High | 59.718 | 10.077 | 18.37 |

In the second step, we used AliSim [35] to simulate DNA sequence alignments along each true gene tree under the General Time-Reversible (GTR) model (using default AliSim GTR model settings) with three different sequence lengths: 400, 100, and 50 bp. In the third step, maximum-likelihood gene trees were inferred using IQ-TREE 2 [41] from the simulated sequence alignments under the Jukes-Cantor (JC) model. We use the simpler JC model when estimating gene trees, instead of the GTR model used to generate the sequences, since this better captures the biases and limitations of applying standard substitution models to real biological sequences when inferring biological gene trees. Thus, from each dataset of true gene trees, we derive 3 additional datasets of estimated gene trees corresponding to the three sequence lengths. The purpose of the second and third steps above is to generate error-prone gene trees that reflect the reconstruction/estimation error present in real gene trees. We found that the estimated gene trees had average normalized Robinson-Foulds (RF) distances [53] (defined below) of 0.08, 0.22, and 0.35 for sequence lengths 400, 100, and 50 bp, respectively, to the corresponding true gene trees. Since some of the methods also make use of gene tree branch lengths, we additionally measured branch length inference error in the estimated medium DTL gene trees. In particular, since the topologies of the true and estimated gene trees can be different, we compared the leaf branch lengths and the path lengths between each pair of leaves using two metrics. First, we used the mean absolute percentage error (MAPE) to measure the error itself, and second, we used the Pearson correlation coefficient (PCC) to measure the correlation between the estimated and inferred lengths. These branch length error statistics are summarized in Supplementary Table S3. For medium DTL datasets and sequence lengths of 400, 100, and 50 bp, path length MAPEs were 6.9%, 15.8%, and 24.1%, respectively. Overall, estimated branch lengths show strong correlation with true lengths. For example, even with 50 bp sequences, we observe PCC over leaf branch lengths and path lengths of 0.824 and 0.833, respectively.

In the fourth and final step, we used Mr.Bayes 3.2.7 [54] to generate the 1000 posterior gene tree samples per gene family required by AleRax. Consistent with the previous step, these trees were inferred by applying Mr.Bayes to the simulated sequence alignments and using the JC model. Following [45], we ran each MCMC chain for 100,000 generations and sampled every 100 generations, using the maximum-likelihood gene trees generated in the previous (third) step above as starting trees. (Note that burn-in is not needed since we start the MCMC chain from the maximum-likelihood tree; this is consistent with how Mr.Bayes is used in [45].) As before, from each dataset of true gene trees, this creates 3 additional datasets of estimated gene trees, with 1000 estimated gene trees per gene family, corresponding to the three sequence lengths.

Table 1 summarizes the specific ranges of parameter values we explored for the number of gene families, DTL rates, and sequence lengths in the core simulated datasets described above. We evaluated all combinations of these parameter values, resulting in a total of 36 core simulated datasets, with each dataset comprising of 10 replicates created using that specific assignment of parameter values. In addition to these core simulated datasets, we also created corresponding simulated datasets with different relative rates of HGT and gene duplication (as described later in the Results section; see Supplementary Table S1 for specific DTL parameters used), and created datasets with 10 and 100 taxa for the runtime and memory usage analysis. The specific commands used to generate the simulated datasets are available in the supplement.

### 2.4 Description of biological datasets

We assembled two previously used biological datasets of different size, composition, and complexity to assess the accuracy and consistency of species trees inferred by AleRax, PhyloGTP, SpeciesRax and ASTRAL-Pro 2, and traditional non-DTL cognizant methods such as MLSA and tANI [22] (Table 3). To examine the effect of extreme divergence and genome complexity variation on species tree inference, we used a dataset composed of 176 Archaea, which was drawn from [18]. The Archaea included in the dataset span 2-3 kingdoms (or superphylums), and radically different lifestyles (from extremophiles inhabiting Antarctic lakes to mammal gut constituents). Because the pan-genome of an entire domain would be immeasurably large and computationally infeasible to accurately infer, we have reduced the number of gene families in this dataset to 282 core genes, which are shared by all members. This also allows direct comparison of the species trees inferred by the four methods to previously calculated phylogenies in [18] which used the same loci. It should be noted that the 282 gene families used in this analysis have been expanded to include all homologs (paralogs, xenologs, etc.) found in each genome, while only orthologs were used in [18].

**Table 3.** Summary of the two biological datasets.

| Dataset | Number of gene families | Potential biases | Previous methods used to infer species tree |
| --- | --- | --- | --- |
| 176 Archaea (domain) | 282 | Extreme divergence, long branch attraction, compositional bias | tANI, MLSA, single gene |
| 44 Frankiales (order) | 8,862 | Low divergence, contamination, genome size difference | tANI, MLSA |

To examine the impact of low sequence divergence on species tree inference, we used a dataset of 44 Frankiales genomes, drawn from [22]. These included taxa are all closely related members of the order Frankiales, and as such the entire pan-genome (8,862 gene families with at least 4 sequences) was used for inference. The order Frankiales are composed of nitrogen-fixing symbionts of pioneer flora [61], and although they demonstrate variation in GC content and genome size these factors were previously shown to not bias phylogenetic inference [22].

**Archaea dataset assembly.** Annotated genomes of 176 Archaea used in [18] were collected. The 282 core gene loci described in [18] were used as amino acid query sequences to search every collected genome, using blastp [9] with default parameters (-evalue was changed to 1e-10). All significant sequence for every loci across all genomes were collected (provided they met a length threshold of 50% in reference to the average gene family sequence size to filter partial sequences). Each gene family was then aligned using mafft-linsi [27] with default parameters. These alignments were used for inferring maximum-likelihood gene trees in IQ-Tree 2 [41], where the best substitution model for each gene family was determined using Bayesian Inference Criterion [26]. The resulting maximum-likelihood gene trees were used as input for ASTRAL-Pro 2, SpeciesRax, and PhyloGTP. To generate the input gene trees for AleRax, we used Mr.Bayes to sample 1000 posterior gene trees under the LG amino-acid model [33] running MCMC chain for 100,000 generations, sampling every 100 generations, and starting each chain with the corresponding maximum-likelihood gene tree for each of the 282 gene families.

**Frankiales dataset assembly.** Annotated proteomes of the 44 Frankiales used in [22] were collected. Protein sequences were clustered into gene families and using the OrthoFinder2.4 pipeline [17] with default parameters (the search algorithm was changed to blast). Briefly, all-vs-all blastp (evalue of 1e-3) was used to find the best hits between input species. The set of query-matches were then clustered into gene families using the MCL algorithm, and the subsequent gene families were aligned using mafft-linsi with default parameters. Resulting alignments were used to create maximum-likelihood gene trees using FastTree [49] using the JTT model and default parameters. To create the input gene trees for AleRax, we used Mr.Bayes to sample 1000 posterior gene trees for each of the 8,862 gene families using the same parameter settings as before.

## 3 Results

### 3.1 Results on simulated data

**Accuracy on true (error-free) gene trees.** We first evaluate the accuracy of the species tree reconstruction methods when given true (error-free) gene trees as input (effectively skipping steps 2, 3 and 4 of the simulation pipeline). While error-free gene trees do not capture the complexities of real data, this analysis helps us understand how the different methods perform in a controlled, ideal setting. Figure 1 shows the results for low, medium, and high DTL rates with varying numbers of gene families for 50-taxon datasets. Unsurprisingly, we find that both DTL rates and number of input gene families are highly impactful parameters. The performance of all four methods worsens as DTL rates increase, and improves as the numbers of input gene families increase. As the figure shows, AleRax shows the highest overall accuracy on these datasets, with PhyloGTP showing comparable but slightly worse accuracy than AleRax. Between PhyloGTP and SpeciesRax, we find that PhyloGTP shows higher accuracy when the number of gene families is small (100 or fewer), particularly when DTL rates are medium or high. For the remaining datasets, both PhyloGTP and SpeciesRax show nearly identical accuracies. Notably, AleRax, PhyloGTP, and SpeciesRax substantially outperform ASTRAL-Pro 2, especially on the medium and high DTL datasets. In particular, we find that ASTRAL-Pro 2 is highly susceptible to high DTL rates, and that it also shows poor performance when the number of input gene families is small. Interestingly, the accuracy of Astral-pro 2 improves rapidly as the number of gene families increases, with the method performing equivalently to the other methods on the low and medium DTL datasets when the input consists of 1000 gene families.

**Figure 1.**
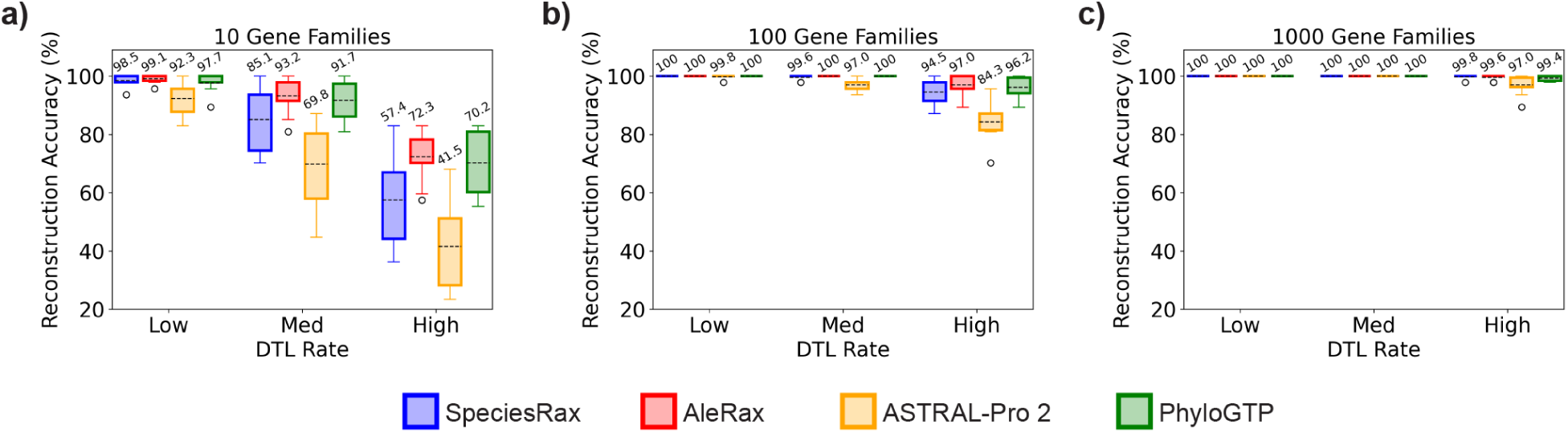
Accuracy on true gene trees. Tree reconstruction accuracies are shown for SpeciesRax, AleRax, ASTRAL-Pro 2, and PhyloGTP when applied to error-free or ‘true’ gene trees. Results are shown for increasing numbers of input gene families (10, 100, and 1000) and for low, medium, and high DTL rates. The number of taxa (i.e., number of leaves in the species tree) is fixed at 50. Higher percentages (y-axis) imply greater accuracy. The number above each box is the mean value across 10 replicate runs, and the dotted line within each box represents the median value.

**Accuracy on estimated (error-prone) gene trees.** We next assess the accuracy of reconstructed species trees when the input consists of estimated (error-prone) gene trees. Figure 2 shows the results of this analysis for all 27 combinations of number of input gene families, DTL rates, and sequence lengths (or gene tree estimation error rates). As expected, the accuracy of all three methods is substantially affected by the quality of the estimated gene trees, with higher accuracies achieved using gene trees estimated from longer sequences. We also find that an increased number of input gene trees can partly make up for error in the input gene trees. AleRax, the only method that explicitly accounts for gene tree inference error and uncertainty, shows the best overall performance, achieving an average reconstruction accuracy of 91.3% when averaged across all 27 datasets with estimated gene trees. PhyloGTP shows the next best accuracy, with an average reconstruction accuracy of 88.8%, and SpeciesRax and ASTRAL-Pro 2 show average accuracies of 86.9% and 81.2%, respectively. As the figure shows, the magnitude of improvement offered by AleRax over the other methods increases as the quality of the input gene trees decreases (i.e., with decreasing sequence length). This is not surprising and points to the significant impact of AleRax’s ability to handle gene tree error and uncertainty. Interestingly, PhyloGTP outperforms AleRax on 6 of the 9 datasets that use the highest quality estimated gene trees (400 base pair sequences; plots in first column of Figure 2). We also find that all methods still outperform ASTRAL-Pro 2 across most datasets and that ASTRAL-Pro 2 continues to be more susceptible to high DTL rates than the other methods. As before, the performance of ASTRAL-Pro 2 improves rapidly with increasing number of input gene trees, even sometimes outperforming all other methods when DTL rates are low or medium. This suggests that ASTRAL-Pro 2 may be well-suited for microbial phylogenomics on datasets with lots of gene trees and relatively low prevalence of HGT. Comparing PhyloGTP with SpeciesRax, we find that both methods have similar performance overall, with PhyloGTP and SpeciesRax showing average percent accuracies of 88.8% and 86.9%, respectively, when averaged across all 27 datasets. However, PhyloGTP shows substantially higher accuracy than SpeciesRax on datasets with high DTL rates, as well as on datasets with 10 input gene trees. This suggests that PhyloGTP may be especially useful for analyzing datasets with high levels of HGT or with a small number of gene trees.

**Figure 2.**
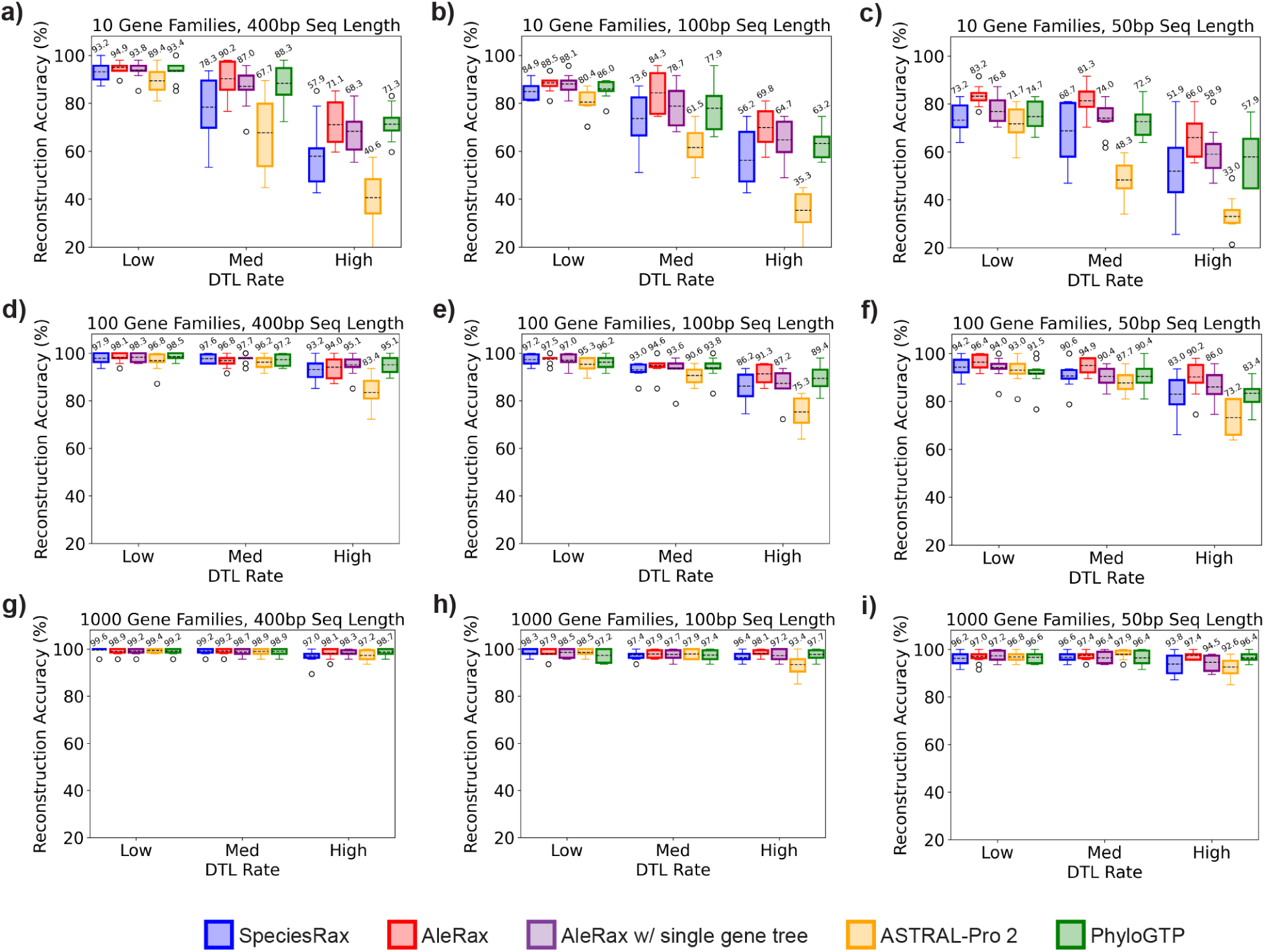
Accuracy on estimated gene trees. Tree reconstruction accuracies are shown for SpeciesRax, AleRax, ASTRAL-Pro 2, and PhyloGTP when applied to estimated gene trees. Results are shown for all 27 combinations of number of input gene families, sequence lengths (shorter sequence lengths imply greater gene tree estimation error), and DTL rates. The first, second, and third rows correspond to datasets with 10, 100, and 1000 gene families, respectively, and the first, second, and third columns correspond to 400, 100, and 50 base pair sequence lengths, respectively. The number of taxa (i.e., number of leaves in the species tree) is fixed at 50. “AleRax” (red) refers to the default execution of AleRax with 1000 gene tree samples per gene family (i.e., with gene tree error-correction), and “AleRax w/single gene tree” (purple) refers to the modified execution where only a single gene tree per gene family is provided as input (i.e., no gene tree error-correction). Higher percentages imply greater accuracy.The number above each box is the mean value across 10 replicate runs, and the dotted line within each box represents the median value.

**Impact of gene tree error-correction on AleRax’s accuracy.** Our results show that AleRax can significantly outperform the other methods on datasets with error-prone gene trees. To better understand how AleRax’s ability to handle gene tree error/uncertainty contributes to its species tree reconstruction accuracy, we also apply AleRax to the estimated (error-prone) gene tree datasets with only a single gene tree per gene family provided as input. Providing only a single gene tree per gene family, instead of the default of 1000, effectively prevents AleRax from being able to account for gene tree error/uncertainty. Figure 2 shows the results of this analysis and reveals several interesting insights (see results for “AleRax w/single gene tree” in that figure). First, we find that the accuracy of this restricted version of AleRax remains quite high and comparable to that of PhyloGTP overall. Second, gene tree error-correction becomes much more impactful on datasets with gene trees of lower quality (100 and 50 base pair sequence lengths). In fact, on the datasets with 400 base pair gene trees, the restricted version of AleRax often slightly outperforms regular AleRax. And third, AleRax’s ability to handle gene tree error/uncertainty is responsible for about a 20% average reduction in reconstruction error on these datasets. These results show that explicit handling of gene tree error and uncertainty can lead to substantially improved species tree reconstruction accuracy, especially on datasets with low-quality gene trees.

**Robustness of PhyloGTP to event costs.** PhyloGTP relies on a parsimony-based DTL reconciliation algorithm [3] to assess the “fit” of input gene trees with candidate species trees. This reconciliation framework relies on user-specified costs for duplication (D), HGT (T), and loss (L) events to compute optimal reconciliations. By default, PhyloGTP uses D-T-L costs of 2-4-1. To assess the robustness of PhyloGTP to different event costs, we apply PhyloGTP variants using five different D-T-L event costs, 2-2-1, 2-3-1, 2-4-1 (default), 2-5-1, and 1-3-1, to the simulated datasets. Supplementary Figures S1 and S2 shows the results of this analysis for true and estimated gene trees, respectively. As the figures show, the performance of PhyloGTP remains robust to the specific event costs used. We also find that no single variant outperforms the others across all, or even most, datasets, and that each of the five variants emerges as the top performer across at least one of the simulated datasets.

**Robustness of results to relative rates of HGT and gene duplication events.** Since biological microbial datasets can have different relative rates of HGT and gene duplication events, we additionally evaluated the methods on simulated datasets with a different ratio of DTL events. In particular, we used higher rates of HGT than in the core datasets (1.5*×*) and near-zero rates of gene duplication. These event rates are based on an analysis of over 7,500 gene families from 103 Aeromonas strains representing 28 different species [50]. As before the loss rate was set to be equal to the gene duplication rate plus the additive HGT rate. As with the core simulated dataset, we used low, medium, and high rates of DTL, different numbers of gene families, and different sequence lengths to obtain 36 alternative simulated datasets, each with 10 replicates. Supplementary Table S1 shows the specific DTL rates and other parameter settings used to generate these alternative datasets.

Results on these alternative datasets are consistent with those reported above for the core simulated datasets. For example, on the alternative datasets with true (error-free) gene trees, we again find that AleRax and PhyloGTP show greatest accuracy and that all methods substantially outperform ASTRAL-Pro 2 on most datasets (Supplementary Figure S3). Likewise, on the alternative datasets with estimated gene trees, AleRax and PhyloGTP continue to be the two best methods, with AleRax, PhyloGTP, SpeciesRax, and ASTRAL-Pro 2 showing average accuracies of 91.5%, 89.0%, 86.2%, and 81.7% across the 27 datasets, respectively (Supplementary Figure S4).

**Runtimes and memory usage.** We compare the runtimes of the four methods when varying the number of taxa (10, 50, and 100) over low, medium, and high DTL rates. In addition, we also evaluate the impact of the number of input gene trees (100 and 1000) using the 50-taxon dataset. These runtimes are shown in Table 4. All methods have parallel implementations and were allocated 12 cores on a 2.1 GHz Intel Xeon processor with 64 GB of RAM. We find that ASTRAL-Pro 2 is, by far, the fastest method, requiring only about 5 seconds on the high-DTL 50-taxon 1000 gene tree datasets and less than 10 seconds on the high-DTL 100-taxon 100 gene tree datasets. SpeciesRax is also extremely fast, requiring only about 60 seconds and 50 seconds, respectively, on those datasets. AleRax and PhyloGTP are much slower than the other two methods, with AleRax requiring over 3 hours and 6 hours, and PhyloGTP requiring about 3.5 hours and 10.5 hours, respectively, on those same datasets. Thus, the improved accuracy provided by AleRax and PhyloGTP comes at the expense of significantly longer running times. It is surprising that PhyloGTP, despite being parsimony based, has the longest runtimes. This is partly due to PhyloGTP’s use of a more extensive SPR-based local search heuristic, while AleRax and SpeciesRax both use a simpler NNI-based local search heuristic. We note that AleRax also requires additional Bayesian analysis runs to generate its input posterior gene tree samples, which add an additional computational burden not accounted for in the reported runtimes. Further research on AleRax and PhyloGTP may lead to faster running times without negatively impacting their accuracies. For example, using fewer posterior gene tree samples (say 100 instead of 1000) per gene family may be sufficient for most analyses. Likewise, it may be possible to speed up the algorithms and local search heuristics implemented in the current prototype version of PhyloGTP.

**Table 4.** Impact of number of taxa and gene trees on running time. Runtimes in seconds are shown for the three methods for datasets with 10, 50, and 100 taxa and low medium, and high rates of DTL. For the 10- and 100-taxon datasets, the number of input gene trees is 100. For 50-taxon datasets, results are shown for both 100 and 1000 gene trees. The runtimes are based on simulated true input gene trees and are averaged over 10 replicate runs. Each method was allocated 12 cores on a 2.1 GHz Intel Xeon processor with 64 GB of RAM.

| Dataset size | DTL rate | SpeciesRax | AleRax | ASTRAL-Pro 2 | PhyloGTP |
| --- | --- | --- | --- | --- | --- |
| 10 taxa,<br>100 gene trees | low | 1.45 | 8.02 | 0.08 | 4.02 |
|  | med | 1.36 | 8.89 | 0.08 | 4.45 |
|  | high | 1.35 | 10.6 | 0.09 | 4.82 |
| 50 taxa,<br>100 gene trees | low | 5.69 | 503.8 | 1.11 | 1,299.56 |
|  | med | 6.25 | 947.45 | 1.14 | 1,374.92 |
|  | high | 8.9 | 1,276.77 | 1.43 | 2,015.03 |
| 50 taxa,<br>1000 gene trees | low | 50.34 | 7,903.59 | 5.38 | 10,011.79 |
|  | med | 52.33 | 9,443.45 | 5.29 | 11,433.19 |
|  | high | 59.95 | 11,376.92 | 5.55 | 13,137.93 |
| 100 taxa,<br>100 gene trees | low | 22.05 | 10,659.92 | 3.61 | 19,871.48 |
|  | med | 28.90 | 14,110.78 | 4.84 | 32,606.87 |
|  | high | 47.19 | 22,091.67 | 7.15 | 38,259.04 |

We also compare the computational memory requirements of the four methods by measuring the peak memory usage during execution. To evaluate the impact of dataset size, memory was profiled on the high DTL rate datasets with 1) 50 taxa and 1000 gene trees and 2) 100 taxa and 100 gene trees. Peak memory usage statistics appear in Supplementary Table S2. We find that PhyloGTP has the lowest memory footprint for the 1000 gene tree dataset while ASTRAL-Pro 2 has the lowest memory footprint for the 100 taxa dataset. Overall, PhyloGTP, ASTRAL-Pro 2, and AleRax have very modest memory requirements in the low hundreds of MBs. SpeciesRax uses substantially more memory than the other methods, but still less than 1 GB on both datasets.

### 3.2 Results on biological data

**Archaeal dataset.** A myriad of controversies surround the phylogeny of Archaea. These controversies include the monophyly of the DPANN superphylum [2, 8, 18, 46, 52], the placement of extreme halophiles [1, 18, 46, 58], and the root of the Archaea [51]. These differences in phylogenetic inference are driven by many factors including, but not limited to compositional bias, long branch attraction, extremely small genomes, numerous HGT events, and biased sampling of metagenome-assembled genomes. Thus, it is interesting to evaluate the performance of the four studied methods in the face of these factors. Using 282 unrooted input gene trees, all four methods inferred Archaeal species trees with inaccuracies with respect to commonly accepted placements of groups in previous analyses. These inaccuracies should be interpreted in the context that for several Archaeal clades (mostly halophiles) there is no consistent, consensus position that is universally accepted amongst Archaeaologists. For example, the monophyly of the DPANN superphylum is considered by some to be an artifact (driven by long branch attraction or biased genome sampling) [2, 18, 72].

There are no extreme topological differences between the SpeciesRax, ASTRAL-Pro 2 and PhyloGTP Archaea tree reconstructions. Those three methods fail to recover a monophyletic Euryarchaea kingdom (Figures 3A and 4A,B) although these resulting topologies (with the Methanomada and Thermococcales on the branches leading to the TACK group) are consistent with trees in previous attempts to find an alternative root of the Archaea [51]. One notable difference is that compositional attraction may have played a larger role in PhyloGTP and ASTRAL-Pro 2, particularly with halophiles. The Haloarchaea were attracted to the Methanonatronarchaeia and were left out of their accepted position within the Methanotecta (Figure 3A) in the PhyloGTP tree. The Methanonatronarchaeia are typically seen as basal to the Methanotecta but have been attracted closer to the other methanogens in the ASTRAL-Pro 2 tree (Figure 4A). Although the Haloarchaea were correctly placed in the SpeciesRax tree (Figure 4B), they are on an extremely long branch. Incorrect placements of the halophiles Nanohaloarchaea, Haloarchaea and Methanonatronarchaeia are often attributed to compositional bias [18]. These halophiles prefer acidic amino acid residues (such as aspartate and glutamate), on account of their survival strategies in hypersaline environments, and these acidified proteomes attract the placement of these groups together in phylogenetic reconstructions.

**Figure 3.**
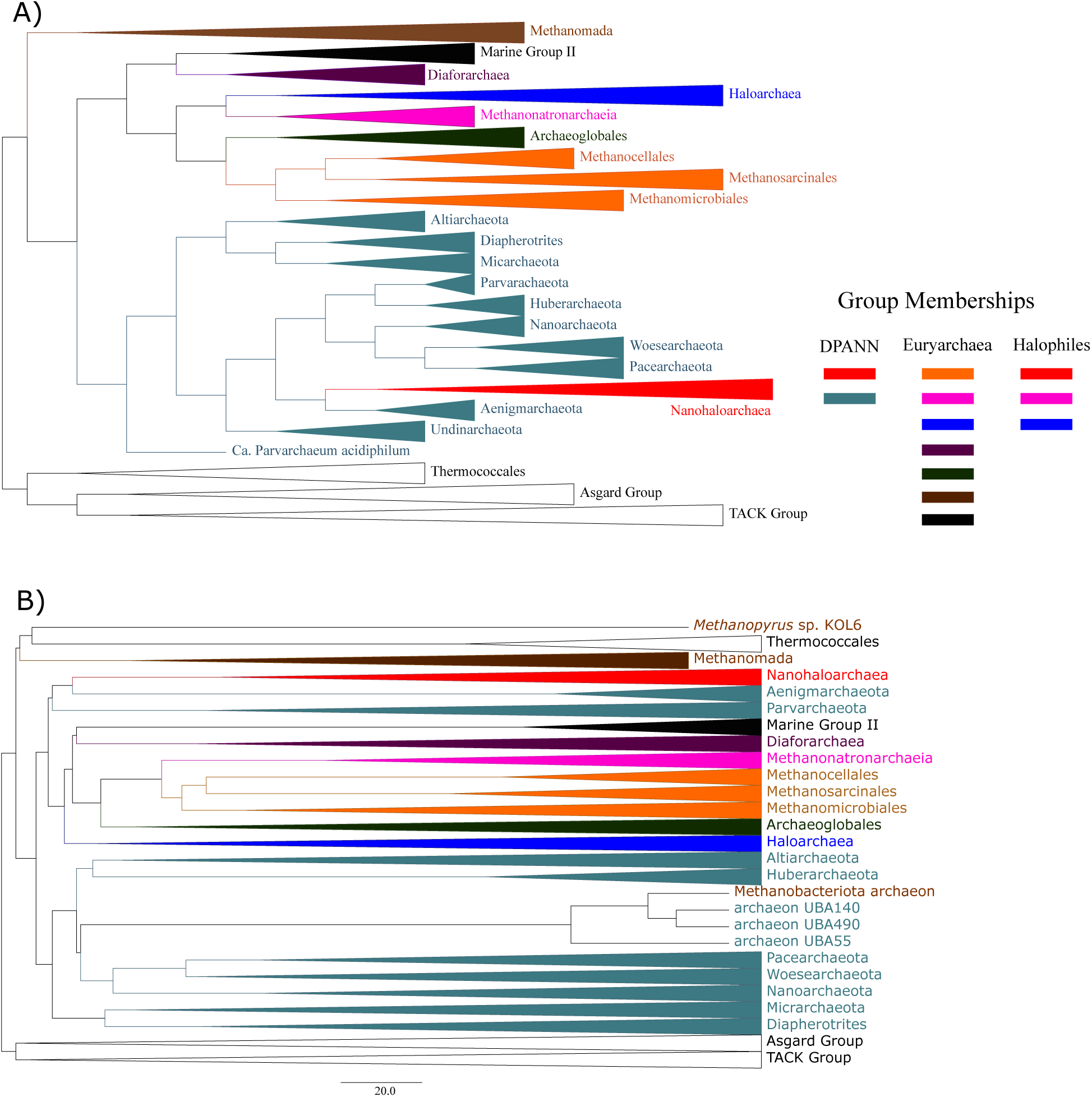
Archaeal species tree reconstructions by PhyloGTP and AleRaX. Individual taxa on both trees have been collapsed into clades and are colored corresponding to higher level classifications (clades with the same color are part of the same class or phylum). The legend shows previously reported Kingdom memberships of these collapsed clades, and also the halophiles which may group together as a result of compositional bias. Part A) Unrooted Archaeal tree inferred by PhyloGTP, shown as a cladogram since PhyloGTP does not infer branch lenghs. Part B) Unrooted Archaeal tree inferred by AleRax.

**Figure 4.**
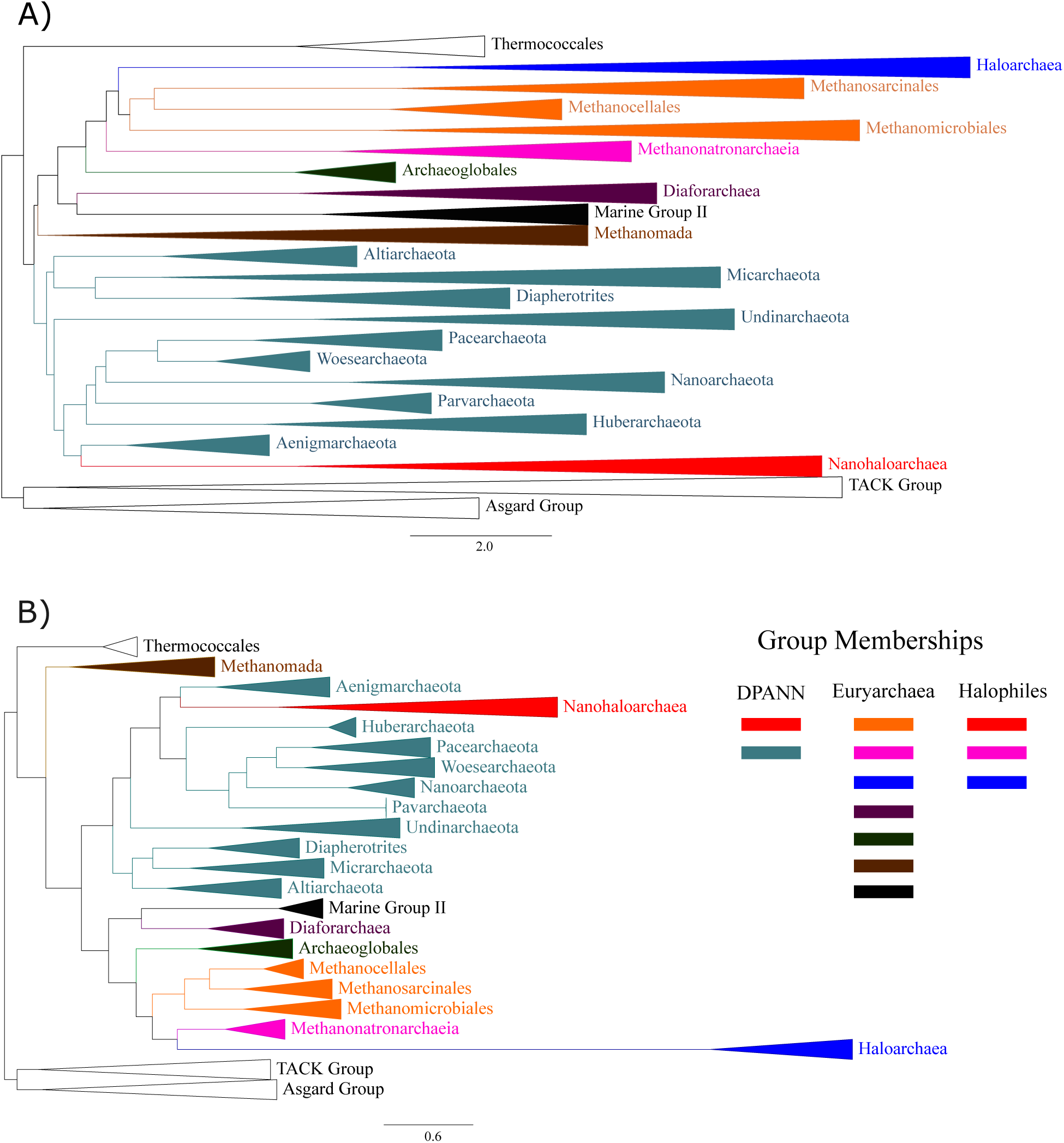
Archaeal species tree reconstructions by ASTRAL-Pro 2 and SpeciesRax. Individual taxa on both trees have been collapsed into clades and are colored corresponding to higher level classifications (clades with the same color are part of the same class or phylum). The legend shows previously reported Kingdom memberships of these collapsed clades, and also the halophiles which may group together as a result of compositional bias. Part A) Unrooted Archaeal tree inferred by ASTRAL-Pro 2. Part B) Unrooted Archaeal tree inferred by SpeciesRax.

In contrast, the AleRax species tree exhibits a markedly different topology (Figure 3B) compared to the above methods.This was the only method that does not recover a monophyletic DPANN group: the Nanohaloarchaea, Parvarchaeota, and Aenigmarchaeota form a clade basal to the Euryarchaea. AleRax fails to recover the correct position of the Haloarchaea, and instead places the group basal to the Methanotecta super-class. Additionally, several Methanomada leaves (*Methanopyrus* sp. Kol6 and *Methanobacteriota archaeaon*) failed to associate with the larger Methanomada clade and were placed in very different places along the tree.

These four different tree topologies for the same input data reveal interesting contrasts in the reconstruction methods. PhyloGTP, SpeciesRax, and ASTRAL-Pro 2 are susceptible to the presence of problematic groups (such as the extreme halophiles) and other biases in complex datasets, potentially limiting their accuracy in some cases. Still, the trees inferred by these three methods are more consistent with previous estimates of the Archaeal tree, demonstrating that these methods can produce a mostly accurate Archaeal tree even in the face of the many biases present in the dataset. In contrast, the tree calculated by AleRax does not resemble any highly supported previously calculated Archaeal tree. This can be explained by two possible causes: 1) AleRax is not suitable for domain level comparisons, where divergence and numerous DTL events have saturated over the extreme time scale (at least 3Ga [39]). 2) The tree produced by AleRax (Figure 3B) reflects the true evolution of the group. Very few previous attempts to consider DTL events in Archaeal phylogenomics exist [14, 66], and were not done at this scale. While these previous studies recovered a monophyletic Euryarchaea and DPANN (in contrast to AleRax), there is not enough information to discount either topology. The topology inferred by AleRax could therefore be viewed as lending credence to the idea that the DPANN are not monophyletic.

**Frankiales dataset.** In the case of the Frankiales, reconstructions with the four methods yield identical relationships between the major clades (Figures 5 and 6). This suggests that all four methods have comparable efficacy when the dataset analyzed is less complex and less divergent. Since this analysis used the entire pan-genome of the Frankiales, a possible concern is that small gene families (such as those that are only found in 4-8 genomes) may negatively impact the methods. To assess the impact of small gene families on species tree reconstruction, a subset of 1,702 genes families present in at least 20 genomes and in the smallest Frankia genome (*Frankia* sp. DG2) was used for inference using PhyloGTP and SpeciesRax. The trees produced from this subset recovered the same topologies for major clades as those in the full complement, indicating that the smaller gene families are not a problem for these methods.

**Figure 5.**
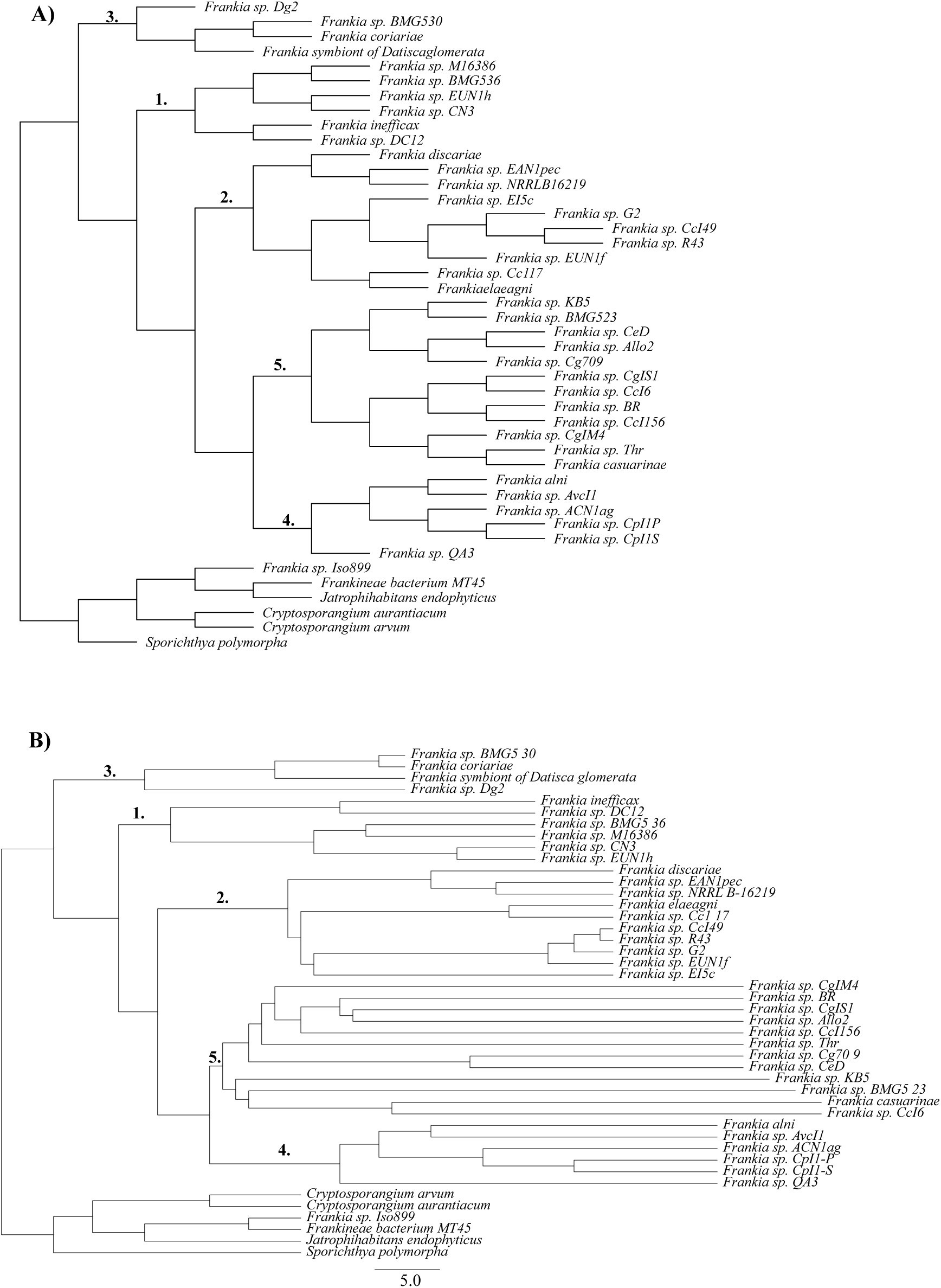
Frankiales species tree reconstructions by PhyloGTP and AleRaX. Clades on both trees are categorized and enumerated based on the group designations described in [22]. Note that both trees show identical relationships among the labeled clades, but not necessarily within those clades. Part A) Unrooted Frankiales cladogram inferred by PhyloGTP. Part B) Unrooted Frankales tree inferred by AleRax.

**Figure 6.**
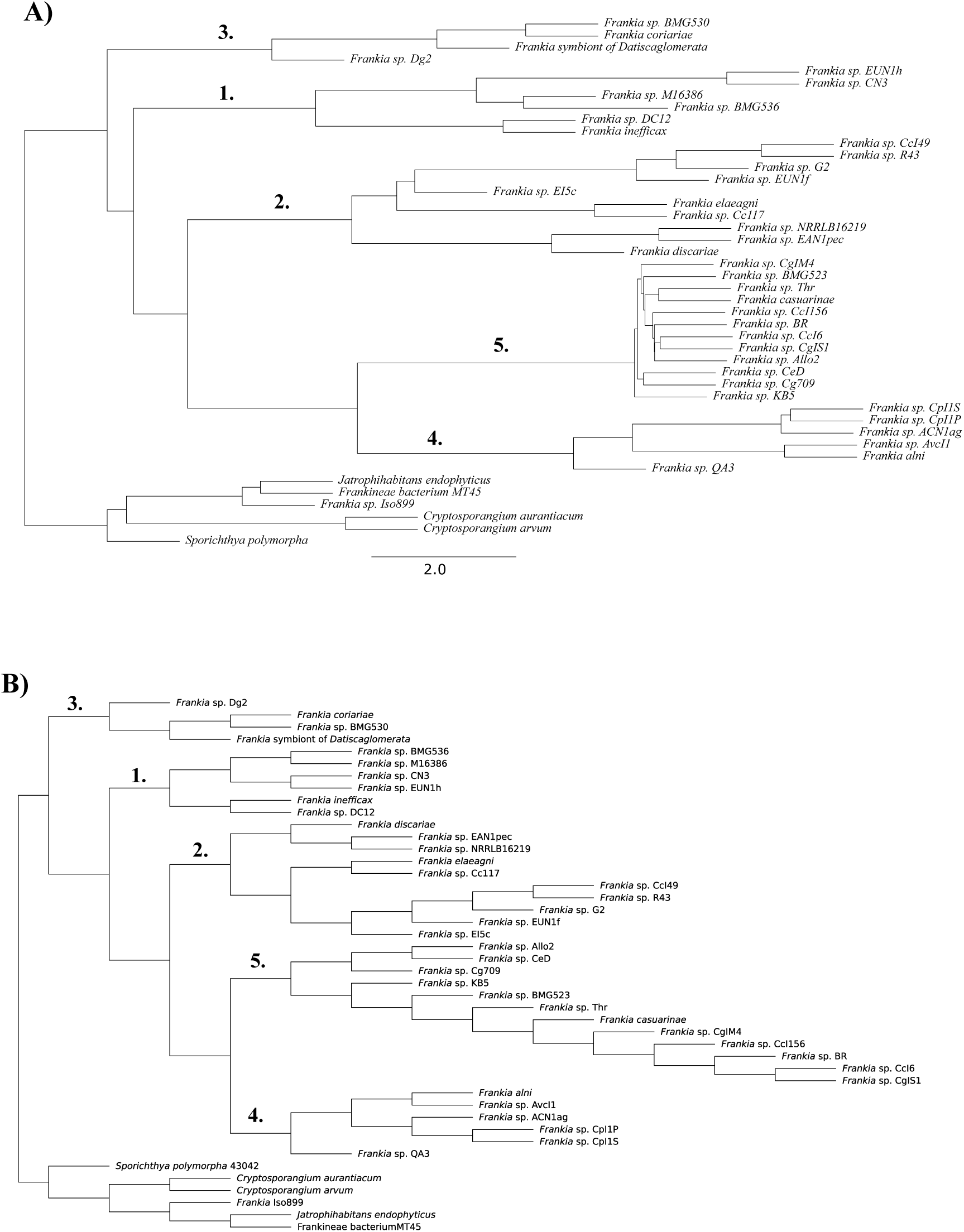
Frankiales species tree reconstructions by ASTRAL-Pro 2 and SpeciesRax. Clades on both trees are categorized and enumerated based on the group designations described in [22]. Note that both trees show identical relationships among the labeled clades, but not necessarily within those clades. Part A) Unrooted Frankiales tree inferred by ASTRAL-Pro 2. Part B) Unrooted Frankales cladogram inferred by SpeciesRax.

In comparison to previous trees inferred on the same genomes using previous methods, such as those shown in [22], there are a few rearrangements of early branching clades in the backbone of the Frankiales. In phylogenies inferred using tANI and MLSA sequence methods, Group 1 (Figure 5) is basal to the rest of the Frankiales. In the trees inferred by the four methods, Group 3 is basal to the other Frankiales, with Group 1 as a later branching basal group. In addition to the movement of these clades, *Frankia* sp. NRRLB16219 and *Frankia* sp. CgIS1 have swapped positions, where *Frankia* sp. CgIS1 has moved from Group 2 to Group 5. These rearrangements may be attributed to the additional genomic data used to reconstruct the four genome-scale trees. Only 24 loci were used in [22], and the inclusion of thousands of additional gene families have painted a slightly different picture of evolution throughout the Frankiales. This suggests that truly genome-scale methods like AleRax, PhyloGTP, ASTRAL-Pro 2, and SpeciesRax could lead to more accurate phylogenomic inference on real datasets compared to other methods. These results also suggest that all four methods can perform well when analyzing less divergent datasets with large numbers of gene families.

## 4 Discussion

In this work, we systematically evaluated four recently developed methods, ASTRAL-Pro 2, SpeciesRax, PhyloGTP, and AleRax, for microbial phylogenomics. These methods can all use thousands of complete (multi-copy) gene families, thereby enabling truly genome-scale microbial phylogenomic species tree inference. Our simulation study identifies AleRax, the only method that explicitly accounts for gene tree inference error/uncertainty, as the best species tree reconstruction accuracy among all tested methods. PhyloGTP shows the best overall accuracy among methods that do not explicitly account for gene tree error and uncertainty, performing particularly well on datasets with high DTL rates or a small number of gene families. Experiments using AleRax with and without gene tree error-correction show that error-correction can lead to an approximately 20% reduction in species tree reconstruction error, especially when the input gene trees are of poor quality.

We also find that AleRax, PhyloGTP, and SpeciesRax almost always outperform ASTRAL-Pro 2, a highly scalable but HGT-naive method. However, our experiments also show that ASTRAL-Pro 2 matches the accuracies of the best methods on datasets with low or medium rates of DTL and a large number of input gene families (1000 in our tests). This suggests that ASTRAL-Pro 2, as well as other closely related quartet based methods such as DISCO-ASTRAL [67], could be potentially useful for analyzing such datasets, especially given their exceptional speed. Another potential advantage of ASTRAL-Pro 2 is that it is agnostic to the underlying evolutionary processes and may therefore be more robust to the effect of evolutionary processes such as incomplete lineage sorting on the datasets being analyzed. On the other hand, the robustness of ASTRAL-Pro 2 to HGT may only hold under simple stochastic models of HGT [13], of the kind employed in our simulation study. ASTRAL-Pro 2 may therefore be more susceptible than other methods to non-random patterns of HGT, such as when HGT frequency depends on the phylogenetic distance between donor and recipient or when some donor-recipient pairs are more likely to engage in HGT than others.

Application of these methods to the two biological datasets of different complexities provides additional valuable insights. We find that all four methods perform well on the less complex Frankiales dataset, but see mixed results on the more complex Archaeal dataset. This disparity was an intentional element of our study design, as these datasets represent opposite ends of a DTL complexity spectrum. The Frankiales dataset, with its higher phylogenetic resolution serves as a transitional case between simulation and highly complex empirical scenarios. Conversely, the Archaeal dataset, with its extensive evolutionary time and documented phylogenetic controversies, represents a stress test for these methods under confounding conditions.

PhyloGTP, SpeciesRax and ASTRAL-Pro 2 produce Archaeal trees that are mostly consistent with current estimates but also have some clearly incorrect placements. On the other hand, AleRax produces a tree that is markedly different than any highly supported previously calculated Archaeal tree. This may be because AleRax is unable to perform well on this complex, highly divergent dataset, or because the AleRax tree more accurately reflects the true evolutionary history of this group. The difficulty in resolving Archaeal phylogenies is well-documented in the literature and stems from a manifold of factors including HGT, compositional biases and long phylogenetic distances. Rather than viewing these inconsistencies as methodological failures, we interpret them as informative indicators of each method’s behavior when confronted with increased evolutionary complexity. Such benchmarking against empirical datasets of varying complexity provides a more realistic assessment of method performance than simulations alone.

Overall, these results suggest that all tested methods are potentially susceptible to compositional and other biases present in complex datasets, and that the results of AleRax, in particular, may need to be interpreted with caution. Our findings underscore the importance of method selection based on expected dataset complexity and highlight the need for careful evaluation of results when analyzing domain-level phylogenies with extensive evolutionary histories. The disparity between simulation and empirical outcomes also suggests opportunities for further methodological refinement.

This work identifies several directions for future research on microbial phylogenomics. First, given our findings on the Archaeal dataset, it would be useful to develop simulation frameworks that better incorporate the specific challenges present in complex empirical datasets. Such advances would help bridge the current gap between theoretical performance and practical application. Second, and related to the above point, it may be informative to perform an expanded simulation study that assesses the impact of additional evolutionary parameters not assessed in the current study. For example, one could assess the impact of heterotachy along species tree branches, distance-dependent (non-uniform) HGT rates, presence of incomplete lineage sorting and gene conversion, compositional biases, etc. Third, the two most accurate methods, AleRax and PhyloGTP, are also the slowest, by far, and could therefore benefit from further methodological and algorithmic development and optimization. And fourth, our results indicate that most methods could benefit by implementing gene tree error-correction or using other strategies to account for gene tree error and uncertainty.

## Acknowledgements

We thank Kashvi Parashar, a high-school student who worked in MSB’s lab during Summer 2024, for helping assess the biological realism of our simulated datasets by comparing their rates of duplication, HGT and loss to those found in real biological datasets.

## Data, script and code availability

Software implementations of all methods evaluated in this work are freely available through their respective web pages or repositories. Scripts used to generate simulated datasets are available in the Supplementary Material. The two real datasets are derived from [18] and [22] and the corresponding alignments and scripts are freely available from Zenodo at the following URL: https://doi.org/10.5281/zenodo.14213472

## References

[1] M. Aouad, G. Borrel, C. Brochier-Armanet, and S. Gribaldo. Evolutionary placement of methanonatronarchaeia. Nature microbiology, 4(4):558–559, 2019.

[2] M. Aouad, N. Taib, A. Oudart, M. Lecocq, M. Gouy, and C. Brochier-Armanet. Extreme halophilic archaea derive from two distinct methanogen class ii lineages. Molecular phylogenetics and evolution, 127:46–54, 2018.

[3] M. S. Bansal, E. J. Alm, and M. Kellis. Efficient algorithms for the reconciliation problem with gene duplication, horizontal transfer and loss. Bioinformatics, 28(12):283–291, 2012.

[4] M. S. Bansal and R. Shamir. A note on the fixed parameter tractability of the gene-duplication problem. IEEE/ACM Trans. Comput. Biology Bioinform., 8(3):848–850, 2011.

[5] M. S. Bansal, Y.-C. Wu, E. J. Alm, and M. Kellis. Improved gene tree error correction in the presence of horizontal gene transfer. Bioinformatics, 31(8):1211–1218, 2015.

[6] R. G. Beiko, T. J. Harlow, and M. A. Ragan. Highways of gene sharing in prokaryotes. Proceedings of the National Academy of Sciences of the United States of America, 102(40):14332–14337, 2005.

[7] M. Bordewich and C. Semple. On the computational complexity of the rooted subtree prune and regraft distance. Annals of Combinatorics, 8:409–423, 2004.

[8] C. Brochier-Armanet, P. Forterre, and S. Gribaldo. Phylogeny and evolution of the archaea: one hundred genomes later. Current opinion in microbiology, 14(3):274–281, 2011.

[9] C. Camacho, G. Coulouris, V. Avagyan, N. Ma, J. Papadopoulos, K. Bealer, and T. L. Madden. Blast+: architecture and applications. BMC bioinformatics, 10:1–9, 2009.

[10] R. Chaudhary, M. S. Bansal, A. Wehe, D. Fernandez-Baca, and O. Eulenstein. iGTP: A software package for large-scale gene tree parsimony analysis. BMC Bioinformatics, 11(1):574, 2010.

[11] F. D. Ciccarelli, T. Doerks, C. von Mering, C. J. Creevey, B. Snel, and P. Bork. Toward automatic reconstruction of a highly resolved tree of life. Science, 311(5765):1283–1287, 2006.

[12] L. A. David and E. J. Alm. Rapid evolutionary innovation during an archaean genetic expansion. Nature, 469:93–96, 2011.

[13] R. Davidson, P. Vachaspati, S. Mirarab, and T. Warnow. Phylogenomic species tree estimation in the presence of incomplete lineage sorting and horizontal gene transfer. BMC Genomics, 16 (Suppl 10):S1, 2015.

[14] A. A. Davín, E. Tannier, T. A. Williams, B. Boussau, V. Daubin, and G. J. Szöllősi. Gene transfers can date the tree of life. Nature ecology & evolution, 2(5):904–909, 2018.

[15] W. F. Doolittle. Phylogenetic classification and the universal tree. Science, 284(5423):2124–2128, 1999.

[16] W. F. Doolittle, Y. Boucher, C. L. Nesbo, C. J. Douady, J. O. Andersson, and A. J. Roger. How big is the iceberg of which organellar genes in nuclear genomes are but the tip? Philosophical Transactions of the Royal Society of London. Series B: Biological Sciences, 358(1429):39–58, 2003.

[17] D. M. Emms and S. Kelly. Orthofinder: phylogenetic orthology inference for comparative genomics. Genome biology, 20:1–14, 2019.

[18] Y. Feng, U. Neri, S. Gosselin, A. S. Louyakis, R. T. Papke, U. Gophna, and J. P. Gogarten. The evolutionary origins of extreme halophilic archaeal lineages. Genome biology and evolution, 13(8):evab166, 2021.

[19] S. R. Gadagkar, M. S. Rosenberg, and S. Kumar. Inferring species phylogenies from multiple genes: Concatenated sequence tree versus consensus gene tree. Journal of Experimental Zoology Part B: Molecular and Developmental Evolution, 304B(1):64–74, 2005.

[20] S. P. Glaeser and P. Kãmpfer. Multilocus sequence analysis (mlsa) in prokaryotic taxonomy. Systematic and Applied Microbiology, 38(4):237–245, 2015. Taxonomy in the age of genomics.

[21] J. P. Gogarten, W. F. Doolittle, and J. G. Lawrence. Prokaryotic evolution in light of gene transfer. Molecular Biology and Evolution, 19(12):2226–2238, 2002.

[22] S. Gosselin, M. S. Fullmer, Y. Feng, and J. P. Gogarten. Improving phylogenies based on average nucleotide identity, incorporating saturation correction and nonparametric bootstrap support. Systematic Biology, 71(2):396–409, 2022.

[23] S. R. Henz, D. H. Huson, A. F. Auch, K. Nieselt-Struwe, and S. C. Schuster. Whole-genome prokaryotic phylogeny. Bioinformatics, 21(10):2329–2335, 05 2004.

[24] E. Hilario and J. P. Gogarten. Horizontal transfer of{ATPase} genes – the tree of life becomes a net of life. Biosystems, 31(2–3):111–119, 1993.

[25] R. P. Hirt, J. M. Logsdon, B. Healy, M. W. Dorey, W. F. Doolittle, and T. M. Embley. Microsporidia are related to fungi: Evidence from the largest subunit of rna polymerase ii and other proteins. Proceedings of the National Academy of Sciences, 96(2):580–585, 1999.

[26] S. Kalyaanamoorthy, B. Q. Minh, T. K. Wong, A. Von Haeseler, and L. S. Jermiin. Modelfinder: fast model selection for accurate phylogenetic estimates. Nature methods, 14(6):587–589, 2017.

[27] K. Katoh and D. M. Standley. Mafft multiple sequence alignment software version 7: improvements in performance and usability. Molecular biology and evolution, 30(4):772–780, 2013.

[28] K. T. Konstantinidis and J. M. Tiedje. Genomic insights that advance the species definition for prokaryotes. Proceedings of the National Academy of Sciences, 102(7):2567–2572, 2005.

[29] S. Kundu and M. S. Bansal. SaGePhy: an improved phylogenetic simulation framework for gene and subgene evolution. Bioinformatics, 02 2019.

[30] M. Lafond and C. Scornavacca. On the weighted quartet consensus problem. Theoretical Computer Science, 769:1–17, 2019.

[31] J. M. Lang, A. E. Darling, and J. A. Eisen. Phylogeny of bacterial and archaeal genomes using conserved genes: Supertrees and supermatrices. PLoS ONE, 8(4):e62510, 04 2013.

[32] P. Lapierre, E. Lasek-Nesselquist, and J. P. Gogarten. The impact of hgt on phylogenomic reconstruction methods. Briefings in Bioinformatics, 15(1):79–90, 2014.

[33] S. Q. Le and O. Gascuel. An Improved General Amino Acid Replacement Matrix. Molecular Biology and Evolution, 25(7):1307–1320, 03 2008.

[34] P. O. Lewis, M.-H. Chen, L. Kuo, L. A. Lewis, K. Fucikova, S. Neupane, Y.-B. Wang, and D. Shi. Estimating bayesian phylogenetic information content. Systematic Biology, 65(6):1009–1023, 2016.

[35] N. Ly-Trong, S. Naser-Khdour, R. Lanfear, and B. Q. Minh. AliSim: A Fast and Versatile Phylogenetic Sequence Simulator for the Genomic Era. Molecular Biology and Evolution, 39(5):msac092, 05 2022.

[36] B. Ma, M. Li, and L. Zhang. From gene trees to species trees. SIAM J. Comput., 30(3):729–752, 2000.

[37] W. P. Maddison and D. Maddison. Mesquite: a modular system for evolutionary analysis. version 2.6. http://mesquiteproject.org, 2009.

[38] V. M. Markowitz, I.-M. A. Chen, K. Palaniappan, K. Chu, E. Szeto, M. Pillay, A. Ratner, J. Huang, T. Woyke, M. Huntemann, I. Anderson, K. Billis, N. Varghese, K. Mavromatis, A. Pati, N. N. Ivanova, and N. C. Kyrpides. Img 4 version of the integrated microbial genomes comparative analysis system. Nucleic Acids Research, 42(D1):D560–D567, 2014.

[39] C. A. Martinez-Gutierrez, J. C. Uyeda, and F. O. Aylward. A timeline of bacterial and archaeal diversification in the ocean. eLife, 12:RP88268, dec 2023.

[40] J. O. McInerney, J. A. Cotton, and D. Pisani. The prokaryotic tree of life: past, presentâ€¦and future? Trends in Ecology & Evolution, 23(5):276 –281, 2008.

[41] B. Q. Minh, H. A. Schmidt, O. Chernomor, D. Schrempf, M. D. Woodhams, A. von Haeseler, and R. Lanfear. IQ-TREE 2: New Models and Efficient Methods for Phylogenetic Inference in the Genomic Era. Molecular Biology and Evolution, 37(5):1530–1534, 02 2020.

[42] S. Mirarab, R. Reaz, M. S. Bayzid, T. Zimmermann, M. S. Swenson, and T. Warnow. Astral: genome-scale coalescent-based species tree estimation. Bioinformatics, 30(17):i541–i548, 08 2014.

[43] B. Morel, A. M. Kozlov, A. Stamatakis, and G. J. Szöllősi. Generax: A tool for species-tree-aware maximum likelihood-based gene family tree inference under gene duplication, transfer, and loss. Molecular Biology and Evolution, 37(9):2763–2774, 06 2020.

[44] B. Morel, P. Schade, S. Lutteropp, T. A. Williams, G. J. Szöllősi, and A. Stamatakis. SpeciesRax: A Tool for Maximum Likelihood Species Tree Inference from Gene Family Trees under Duplication, Transfer, and Loss. Molecular Biology and Evolution, 39(2):msab365, 01 2022.

[45] B. Morel, T. A. Williams, A. Stamatakis, and G. J. Szollosi. AleRax: a tool for gene and species tree co-estimation and reconciliation under a probabilistic model of gene duplication, transfer, and loss. Bioinformatics, 40(4):btae162, 2024.

[46] P. Narasingarao, S. Podell, J. A. Ugalde, C. Brochier-Armanet, J. B. Emerson, J. J. Brocks, K. B. Heidelberg, J. F. Banfield, and E. E. Allen. De novo metagenomic assembly reveals abundant novel major lineage of archaea in hypersaline microbial communities. The ISME journal, 6(1):81–93, 2012.

[47] G. J. Olsen, C. R. Woese, and R. Overbeek. The winds of (evolutionary) change: breathing new life into microbiology. Journal of Bacteriology, 176(1):1–6, 1994.

[48] R. D. M. Page. GeneTree: comparing gene and species phylogenies using reconciled trees. Bioinformatics (Oxford, England), 14(9):819–820, 1998.

[49] M. N. Price, P. S. Dehal, and A. P. Arkin. Fasttree 2–approximately maximum-likelihood trees for large alignments. PloS one, 5(3):e9490, 2010.

[50] L. T. Rangel, J. Marden, S. Colston, J. C. Setubal, J. Graf, and J. P. Gogarten. Identification and characterization of putative aeromonas spp. t3ss effectors”. PLOS ONE, 14(6):1–20, 06 2019.

[51] K. Raymann, C. Brochier-Armanet, and S. Gribaldo. The two-domain tree of life is linked to a new root for the archaea. Proceedings of the National Academy of Sciences, 112(21):6670–6675, 2015.

[52] K. Raymann, P. Forterre, C. Brochier-Armanet, and S. Gribaldo. Global phylogenomic analysis disentangles the complex evolutionary history of dna replication in archaea. Genome biology and evolution, 6(1):192–212, 2014.

[53] D. Robinson and L. Foulds. Comparison of phylogenetic trees. Mathematical Biosciences, 53(1):131–147, 1981.

[54] F. Ronquist, M. Teslenko, P. van der Mark, D. L. Ayres, A. Darling, S. Höhna, B. Larget, L. Liu, M. A. Suchard, and J. P. Huelsenbeck. MrBayes 3.2: Efficient Bayesian Phylogenetic Inference and Model Choice Across a Large Model Space. Systematic Biology, 61(3):539–542, 2012.

[55] G. Sevillya, D. Doerr, Y. Lerner, J. Stoye, M. Steel, and S. Snir. Horizontal Gene Transfer Phylogenetics: A Random Walk Approach. Molecular Biology and Evolution, 37(5):1470–1479, 12 2019.

[56] A. Shifman, N. Ninyo, U. Gophna, and S. Snir. Phylo si: a new genome-wide approach for prokaryotic phylogeny. Nucleic Acids Research, 42(4):2391–2404, 2014.

[57] Y. S. Song. On the combinatorics of rooted binary phylogenetic trees. Annals of Combinatorics, 7(3):365–379, 2003.

[58] D. Y. Sorokin, K. S. Makarova, B. Abbas, M. Ferrer, P. N. Golyshin, E. A. Galinski, S. Ciorda, M. C. Mena, A. Y. Merkel, Y. I. Wolf, et al. Reply to ‘evolutionary placement of methanonatronarchaeia’. Nature Microbiology, 4(4):560–561, 2019.

[59] G. J. Szollosi, E. Tannier, N. Lartillot, and V. Daubin. Lateral gene transfer from the dead. Systematic Biology, 62(3):386–397, 2013.

[60] A. Tofigh, M. T. Hallett, and J. Lagergren. Simultaneous identification of duplications and lateral gene transfers. IEEE/ACM Trans. Comput. Biology Bioinform., 8(2):517–535, 2011.

[61] M. Trujillo, S. Dedysh, P. DeVos, B. Hedlund, P. Kampfer, F. Rainey, and W. Whitman. Bergey’s manual of systematics of archaea and bacteria. Wiley Online Library, 2021.

[62] A. Wehe, M. S. Bansal, J. G. Burleigh, and O. Eulenstein. Duptree: a program for large-scale phylogenetic analyses using gene tree parsimony. Bioinformatics, 24(13), 2008.

[63] A. Wehe and J. Burleigh. Scaling the gene duplication problem towards the tree of life. In 2nd International Conference on Bioinformatics and Computational Biology 2010, BICoB 2010, pages 133–138, 01 2010.

[64] S. Weiner, Y. Feng, J. P. Gogarten, and M. S. Bansal. Assessing the potential of gene tree parsimony for microbial phylogenomics. In C. Scornavacca and M. Hernández-Rosales, editors, Comparative Genomics, pages 129–149, Cham, 2024. Springer Nature Switzerland.

[65] C. Whidden, N. Zeh, and R. G. Beiko. Supertrees based on the subtree prune-and-regraft distance. Systematic Biology, 2014.

[66] T. A. Williams, G. J. Szollosi, A. Spang, P. G. Foster, S. E. Heaps, B. Boussau, T. J. G. Ettema, and T. M. Embley. Integrative modeling of gene and genome evolution roots the archaeal tree of life. Proceedings of the National Academy of Sciences, 114(23):E4602–E4611, 2017.

[67] J. Willson, M. S. Roddur, B. Liu, P. Zaharias, and T. Warnow. DISCO: Species Tree Inference using Multicopy Gene Family Tree Decomposition. Systematic Biology, 71(3):610–629, 08 2021.

[68] C. R. Woese. Bacterial evolution. Microbiological Reviews, 51(2):221–271, 1987.

[69] W. H. Yap, Z. Zhang, and Y. Wang. Distinct types of rrna operons exist in the genome of the actinomycete thermomonospora chromogena and evidence for horizontal transfer of an entire rrna operon. Journal of Bacteriology, 181(17):5201–5209, 1999.

[70] C. Zhang and S. Mirarab. ASTRAL-Pro 2: ultrafast species tree reconstruction from multi-copy gene family trees. Bioinformatics, 38(21):4949–4950, 09 2022.

[71] O. Zhaxybayeva, W. F. Doolittle, R. T. Papke, and J. P. Gogarten. Intertwined evolutionary histories of marine synechococcus and prochlorococcus marinus. Genome Biology and Evolution, 1:325–339, 2009.

[72] O. Zhaxybayeva, R. Stepanauskas, N. R. Mohan, and R. T. Papke. Cell sorting analysis of geographically separated hypersaline environments. Extremophiles, 17:265–275, 2013.

