## Supplementary text, tables, and figures for "A systematic assessment of phylogenomic approaches for microbial species tree reconstruction"

### Supplementary Material

### PhyloGTP: Methodological Details

The local search heuristic implemented in PhyloGTP is similar to those use for many other NP-hard phylogeny inference problems, including those used for other popular variants of gene tree parsimony [10, 37, 48, 62]. The local search heuristic starts with an initial candidate rooted species tree and iteratively improves it using local search. Specifically, in each local search iteration, the heuristic finds a minimum reconciliation cost tree in the "local neighborhood" of the current species tree. The best tree found in that local neighborhood then becomes the starting point for the next local search iteration. The heuristic terminates when a lower cost tree cannot be found in the local neighborhood of the current species tree. Next, we describe how PhyloGTP computes the initial candidate species tree and how it defines the local neighborhood for each subsequent local search iteration.

**Construction of initial candidate species tree.** If an estimated user-defined initial species tree is unavailable, PhyloGTP uses a stepwise taxon-addition algorithm to compute a reasonable initial species tree for the local search. The stepwise taxon-addition algorithm works by starting from a two-taxon rooted species tree and iteratively placing taxa, one at a time, onto the species tree topology along the branch that minimizes the total DTL reconciliation cost. In our implementation, the taxa are added in order of decreasing coverage, where the coverage of taxon $s$ is the number of gene trees that include a gene from $s$. At each iteration, each gene tree is pruned to reflect only the taxa present in the current (incomplete) species tree. Once all taxa have been added, the resulting rooted species tree is used as the starting species tree for the subsequent local search. We found that using this stepwise taxon-addition algorithm results in an average reduction of $93\%$ in the number of local search iterations until convergence when compared to using a random species tree topology as the initial starting tree (detailed results not shown).

**Description of local search iterations.** PhyloGTP implements a constrained (rooted) subtree prune and regraft (SPR) [7] based local search using the initial tree as a starting point. SPR is the most commonly used tree edit operation for phylogenetic local search and induces a local neighborhood of $\Theta(n^2)$ trees, where $n$ is the number of leaves in the species tree [57]. Rather than always evaluating all trees in the full SPR neighborhood at each iteration, PhyloGTP first considers only the restricted set of trees obtained by regrafting a single pruned subtree $S_v$, rooted at a some node $v \in V(S)/\,rt(S)$, onto each possible edge in the current species tree $S$. It finds the lowest cost tree $S'$ within that restricted neighborhood and, if $S'$ has lower cost than $S$, then $S$ is replaced by $S'$ and PhyloGTP proceeds to the next local search iteration. If no improvement was found in the restricted neighborhood using $S_v$, then a new node $u \neq v \in V(S)$ is chosen and the restricted local search step is repeated using the pruned subtree $S_u$. Thus, PhyloGTP is initially constrained to a small subset of the full SPR search space, but will incrementally expand the set of trees under consideration until an improvement is found, or until the full SPR neighborhood is explored. In the latter case, if no improvement is found then the search is determined to have converged. Note that the order in which subtrees are considered for pruning is randomized at the beginning of each local search iteration. In addition, if there are multiple species trees with minimum reconciliation cost within a restricted neighborhood, then the new species tree $S'$ is selected uniformly at random among them.

Observe that PhyloGTP use a search strategy based on restricted SPR local neighborhoods instead of exploring the full SPR local neighborhood at each local search iteration. This is motivated by the underlying computational complexity of the computation. If $n$ denotes the number of taxa in the analysis and $k$ the number of input gene trees then, assuming most of the $k$ gene trees have $\Theta(n)$ leaves, the time complexity of naively evaluating all candidate species trees in a single SPR local neighborhood becomes $\Theta(n^2) \times \Theta(n^2) \times \Theta(k)$ which is $\Theta(k \cdot n^4)$. This does not scale well with increasing $n$. Furthermore, many local search iterations have to be performed during a single execution of the heuristic. By using a search strategy based on restricted SPR local neighborhoods, the number of candidate species trees evaluated during most local search iterations reduces to $\Theta(n)$, reducing the time complexity of most local search iterations to a more reasonable $\Theta(k \cdot n^3)$. Importantly, this approach retains the key advantage of using a full SPR-based search since the heuristic search only terminates if a better tree is not found in the full SPR local neighborhood. Previous work on a simpler GTP problem suggests that heuristics based on restricted SPR local neighborhoods can perform almost as well as those based on using full SPR neighborhoods during each local search iteration [63].

**DTL event costs assignment.** By default, PhyloGTP uses event costs of 2, 4, and 1 for gene duplications, HGTs, and gene losses, respectively (i.e., $P_d = 2$, $P_t = 4$, and $P_l = 1$). Unless otherwise noted, experimental results reported in this manuscript are based on these default event costs for PhyloGTP.

**Parallelization.** PhyloGTP implements parallelization to further improve its scalability and enable application to large-scale datasets. The parallelization strategy works by dynamically distributing the computation associated with obtaining the reconciliation costs of candidate species trees in the local search neighborhood across a user-defined number of cores. Thus, when using $c$ cores, the running time of the heuristic is reduced by roughly a factor of $c$.

**Table S1. Key parameters used in the simulation study for the alternative datasets.** The table lists the parameters and their values explored in the simulation study for the alternative simulated datasets with different relative DTL rates. All 36 ($= 3 \times 3 \times 4$) combinations of these three parameters were evaluated at 10 replicates each. DTL rates are specified in the form $(d, t, l)$, where $d$ is the gene duplication rate, $t$ is the HGT rate (split evenly between additive and replacing HGTs), and $l$ is the gene loss rate. The number of species was fixed at 50 for these datasets.

| **Parameter** | **Values** |
|---|---|
| Number of gene families | $10, 100, 1000$ |
| DTL rates | low = $(0.03, 0.9, 0.48)$<br>med = $(0.06, 1.8, 0.96)$<br>high = $(0.12, 3.6, 1.92)$ |
| Sequence length (nucleotides) | $400, 100, 50$, and true gene trees |

**Table S2. Peak memory usage.** Peak memory usage was measured in MB for each method over two datasets with different characteristics. The first row corresponds to datasets with 50 taxa and 1000 gene trees and the second row corresponds to datasets with 100 taxa and 100 gene trees. Results are averaged over the 10 replicate runs in each dataset.

| Dataset Size | SpeciesRax | AleRax | ASTRAL-Pro 2 | PhyloGTP |
|---|---|---|---|---|
| 50 taxa,<br>1000 gene trees | 913.9 | 144.5 | 144.2 | 125.1 |
| 100 taxa,<br>100 gene trees | 298.9 | 84.3 | 27.1 | 81.0 |

**Table S3. Gene tree branch length estimation error metrics.** Error in branch lengths in the estimated gene trees was quantified using two metrics: mean absolute percentage error (MAPE) and Pearson correlation coefficient (PCC). Since the topologies of the true and estimated gene trees are different, we applied these metrics to two measurements: the branch lengths of edges directly above leaf nodes (denoted 'Leaf MAPE' and 'Leaf PCC') and the path lengths between each pair of leaves (denoted 'Path MAPE' and 'Path PCC'). Results are shown on the medium DTL rate datasets over the 3 different sequence lengths of 400, 100, and 50 bp.

| Sequence length | Leaf MAPE | Leaf PCC | Path MAPE | Path PCC |
|---|---|---|---|---|
| 400 | 0.253 | 0.976 | 0.069 | 0.982 |
| 100 | 0.493 | 0.902 | 0.158 | 0.909 |
| 50 | 0.666 | 0.824 | 0.241 | 0.833 |

#### Supplementary figures

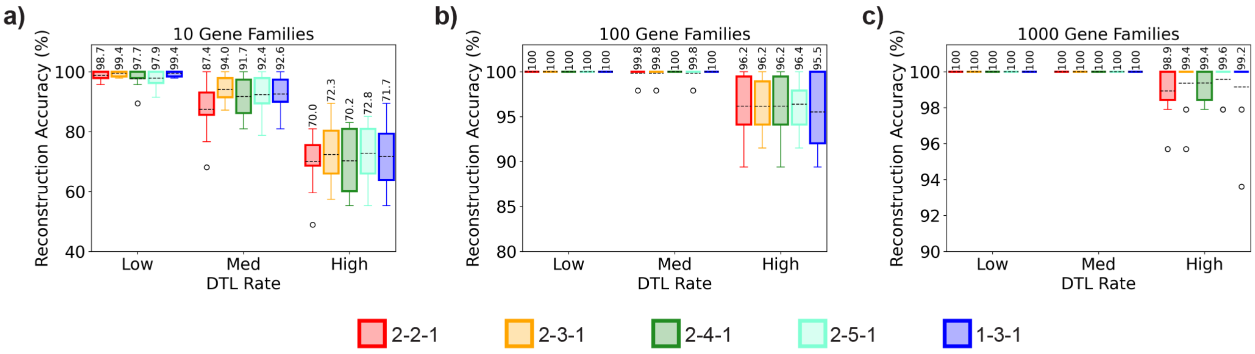


**Figure S1. Impact of DTL event costs on PhyloGTP's accuracy on true gene trees.** Tree reconstruction accuracies are shown for five variants of PhyloGTP using different costs for duplication (D), transfer (T), and loss (L) events when applied to true (error-free) gene trees. The event cost settings used by each variant are specified in the order D-T-L. Setting 2-4-1 correspond to the default version of PhyloGTP. Results are shown for increasing numbers of input gene families (10, 100, and 1000) and for low, medium, and high DTL rates. The number of taxa (i.e., number of leaves in the species tree) is fixed at 50. Higher percentages (y-axis) imply greater accuracy. The number above each box is the mean value across 10 replicate runs, and the dotted line within each box represents the median value.

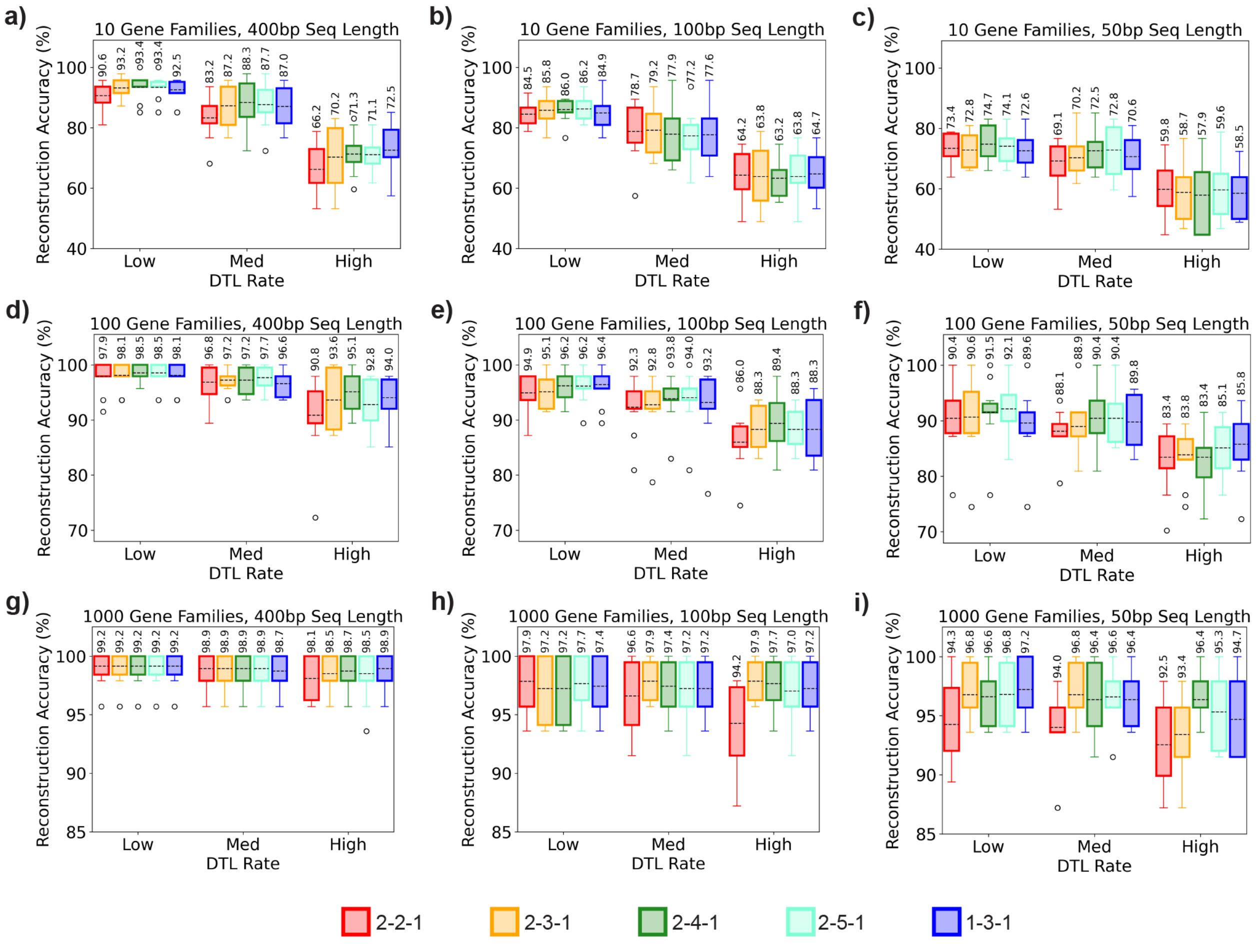


**Figure S2. Impact of DTL event costs on PhyloGTP's accuracy on estimated gene trees.** Tree reconstruction accuracies are shown for five variants of PhyloGTP using different costs for duplication (D), transfer (T), and loss (L) events when applied to estimated (error-prone) gene trees. The event cost settings used by each variant are specified in the order D-T-L. Setting 2-4-1 correspond to the default version of PhyloGTP. Results are shown for all 27 combinations of number of input gene families, sequence lengths (shorter sequence lengths imply greater gene tree estimation error), and DTL rates. The first, second, and third rows correspond to datasets with 10, 100, and 1000 gene families, respectively, and the first, second, and third columns correspond to 400, 100, and 50 base pair sequence lengths, respectively. The number of taxa (i.e., number of leaves in the species tree) is fixed at 50. Higher percentages (y-axis) imply greater accuracy. The number above each box is the mean value across 10 replicate runs, and the dotted line within each box represents the median value.

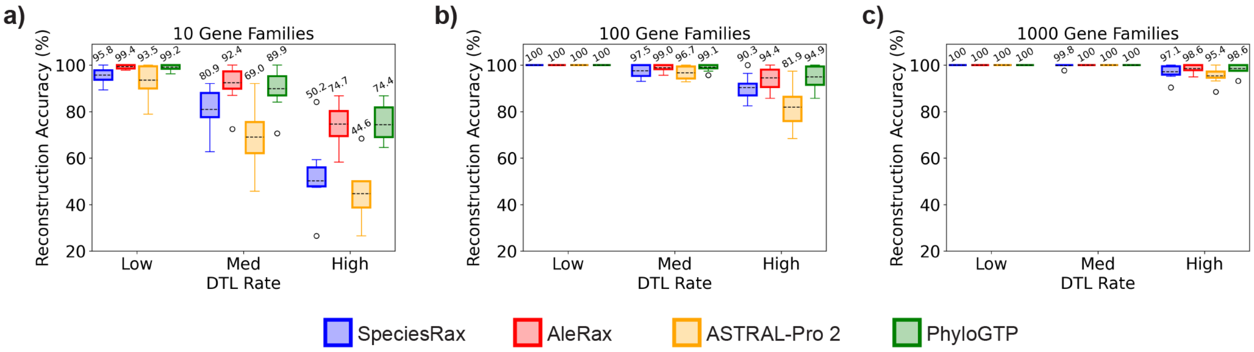


**Figure S3. Accuracy with true gene trees for the alternative simulated datasets.** Tree reconstruction accuracies are shown for SpeciesRax, AleRax, ASTRAL-Pro 2, and PhyloGTP when applied to error-free or 'true' gene trees on the alternative datasets with different relative DTL rates. Results are shown for increasing numbers of input gene families (10, 100, and 1000) and for low, medium, and high DTL rates. The number of taxa (i.e., number of leaves in the species tree) is fixed at 50. Higher percentages (y-axis) imply greater accuracy. The number above each box is the mean value across 10 replicate runs, and the dotted line within each box represents the median value.

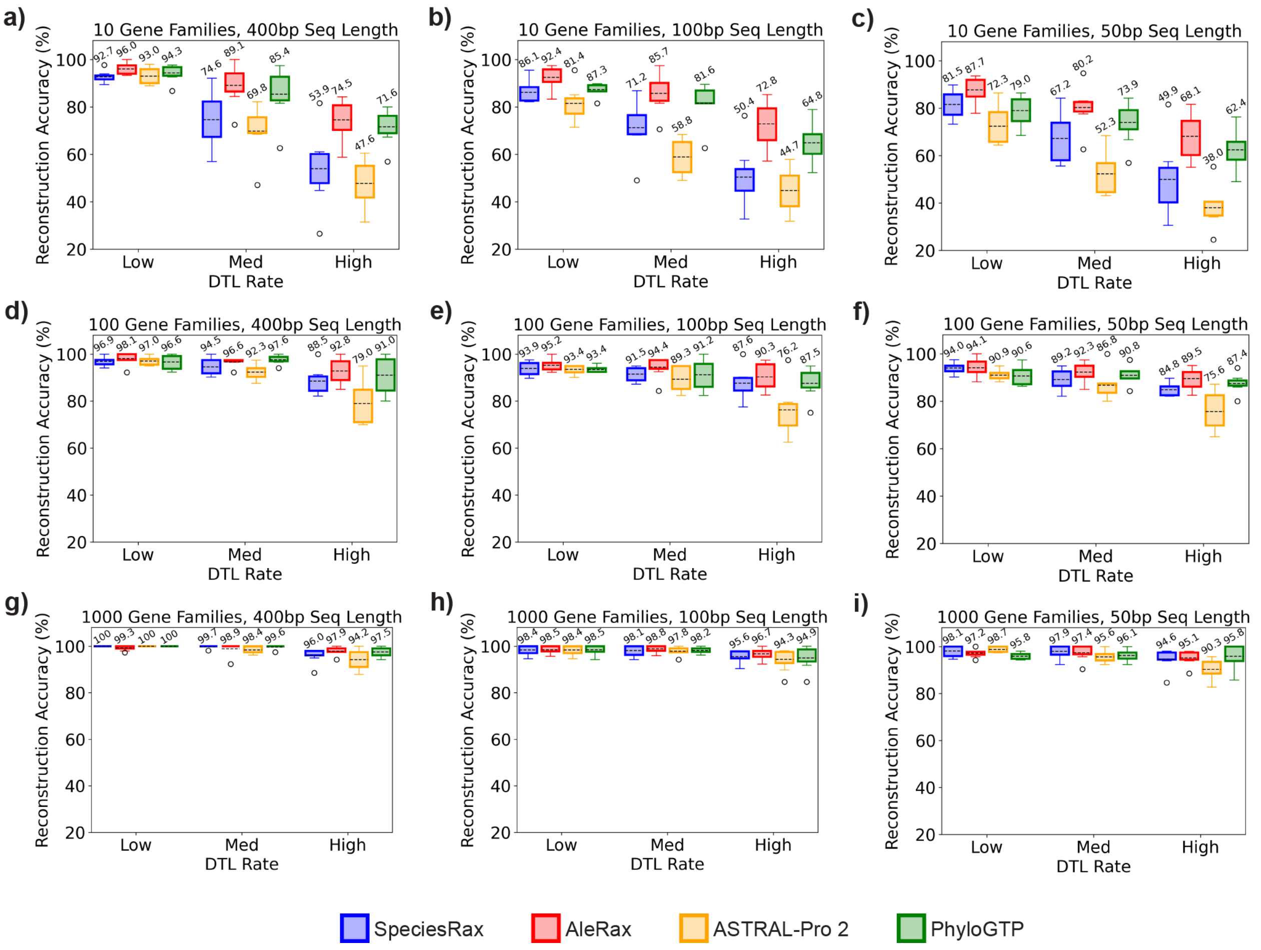


**Figure S4. Accuracy with estimated gene trees for the alternative simulated datasets.** Tree reconstruction accuracies are shown for SpeciesRax, AleRax, ASTRAL-Pro 2, and PhyloGTP when applied to estimated gene trees on the alternative datasets with different relative DTL rates. Results are shown for all 27 combinations of number of input gene families, sequence lengths (shorter sequence lengths imply greater gene tree estimation error), and DTL rates. The first, second, and third rows correspond to datasets with 10, 100, and 1000 gene families, respectively, and the first, second, and third columns correspond to 400, 100, and 50 base pair sequence lengths, respectively. The number of taxa (i.e., number of leaves in the species tree) is fixed at 50. Higher percentages imply greater accuracy. The number above each box is the mean value across 10 replicate runs, and the dotted line within each box represents the median value.

#### Scripts for generating simulated datasets

This section provides the scripts which can be used to recreate the simulated datasets used in this study. The first script creates the true species tree and error-free gene trees with SaGePhy [29]. Note that the variables `n_leaf`, `n_genes`, `d`, `l`, and `t` should be changed as needed to generate the full spectrum of datasets (see, e.g., Table 1 in main manuscript and Supplementary Table S1).

```
#!/bin/bash

sage_path=sagephy/sagephy-1.0.0.jar
out_path=out_true
n_leaf=50
n_genes=100
birth=5.0
death=2.5

d=0.3
l=0.6
t=0.6

mkdir $out_path
java -jar $sage_path HostTreeGen -nox -min $n_leaf -max $n_leaf -a 100000 1.0 $birth
    $death ${out_path}/spec

mkdir ${out_path}/gene_trees
for ((i=1;i<=$n_genes;i++)); do
    java -jar $sage_path GuestTreeGen -min 10 -nox  ${out_path}/spec.pruned.tree $d $l
     $t ${out_path}/gene_trees/gene.${i}
done
```

The second script generates sequences for each of the gene trees generated in the previous step using AliSim [35], then computes ML error-prone gene trees with IQ-TREE 2 [41]. Note that the variables `n_genes` and `seq_length` and paths to input gene trees should be changed as needed to generate the full spectrum of datasets.

```
#!/bin/bash

n_genes=100
seq_length=400

gene_tree_dir=out_true/gene_trees
out_dir=out_inferred/seqs

mkdir $out_path
for ((i=1;i<=$n_genes;i++)); do
    tree_path=${inp_path}/gene_trees/gene.${i}.pruned.tree
    out_path=${out_dir}/gene.${i}
    iqtree2 --alisim ${out_path} -m GTR -t $tree_path --length $seq_length
    iqtree2 -s ${out_path}.phy -m JC --prefix ${out_path}.phy
done
```

We also provide a MrBayes [54] script which can be used to generate the input posterior gene tree samples for evaluating AleRax from the sequences generated with the previous script.

```
begin mrbayes;
    set autoclose=yes nowarn=yes;
```

```
    execute nex_path;
    startvals tau=usertree;
    lset nst=1;
    prset statefreqpr=fixed(equal);
    mcmc nchains=2 nruns=1 ngen=100000 samplefreq=100 Checkpoint=No Stoprule=No;
end;
```
